# The interplay between molten globules and heme disassociation defines human hemoglobin disassembly

**DOI:** 10.1101/726216

**Authors:** P.P. Samuel, M.A. White, W.C. Ou, D.A. Case, G.N. Phillips, J.S. Olson

## Abstract

Hemoglobin functions as an oxygen transport protein, with each subunit containing a heme cofactor. We have developed a global disassembly model for human hemoglobin, linking hemin (ferric heme) disassociation and apo(heme-free)-protein unfolding pathways. The model was based on the evaluation of circular dichroism and visible absorbance measurements of guanidine hydrochloride-induced disassembly of holo (heme-bound)-hemoglobin and previous measurements of apohemoglobin unfolding. The populations of holo-intermediates and equilibrium disassembly parameters were determined quantitatively for adult and fetal hemoglobins. The key stages for disassembly into unfolded monomers are characterized by hemichrome intermediates with molten globule characteristics. Hemichromes, which occur when both hemin iron axial sites coordinate amino acids, are not energetically favored in native human hemoglobins. However, these hexacoordinate iron complexes are important for preventing hemin disassociation from partially unfolded species during early disassembly and late stage assembly events. Both our model evaluation and independent small angle X-ray scattering measurements demonstrate that heme disassociation during early disassembly leads to loss of tetrameric structural integrity. Dimeric and monomeric hemichrome intermediates occur along the disassembly pathway inside red cells where the hemoglobin concentration is very high. This prediction explains why in the red cells of patients with unstable hemoglobinopathies, misassembled hemoglobins often get trapped as hemichromes that accumulate into insoluble Heinz bodies. These Heinz bodies become deposited on the cell membranes and can lead to hemolysis. Alternatively, when acellular hemoglobin is diluted into blood plasma after red cell lysis, the disassembly pathway is dominated by early hemin disassociation events, which leads to the generation of higher fractions of apo-subunits and free hemin known to damage to the integrity of blood vessel walls. Thus, our model illuminates the pathophysiology of hemoglobinopathies and other disease states associated with unstable globins and red cell lysis, and provides insights into the factors governing hemoglobin assembly during erythropoiesis.

**Significance:** Our deconvolution and global analysis of spectral data led to both the characterization of “hidden” hemichrome intermediates and the development of a quantitative model for human hemoglobin disassembly/assembly. The importance of this mechanism is several-fold. First, the hemoglobin system serves as a general biological model for understanding the role of oligomerization and cofactor binding in facilitating protein folding and assembly. Second, the fitted parameters provide: (a) estimates of hemin affinity for apoprotein states; (b) quantitative interpretations of the pathophysiology of hemoglobinopathies and other diseases associated with unstable globins and red cell lysis; (c) insights into the factors governing hemoglobin assembly during erythropoiesis; and (d) a framework for designing targeted hemoglobinopathy therapeutics.

## Introduction

An interwoven relationship between protein function, structure, and assembly occurs during hemoglobin assembly through step-wise hemin (Fe(II)-protoporphyrin IX) binding that guides the formation of secondary, tertiary, and quaternary protein structures (Fig. 1). We depend on erythrocyte-encapsulated hemoglobin (Hb) to transport oxygen in our cardiovascular system (1). Hb regulates cooperative binding of oxygen to each of its ferrous heme iron atoms via its quaternary conformations. Functional adult HbA consists of α and β globin subunits interacting at both the α_1_β_1_ dimer and α_1_β_2_ tetramer interfaces. Each globin subunit has 7 or 8 alpha helical segments labeled A to H, and is folded with a heme molecule sandwiched within a cavity between the E and F helices (2, 3).

**Figure 1.**
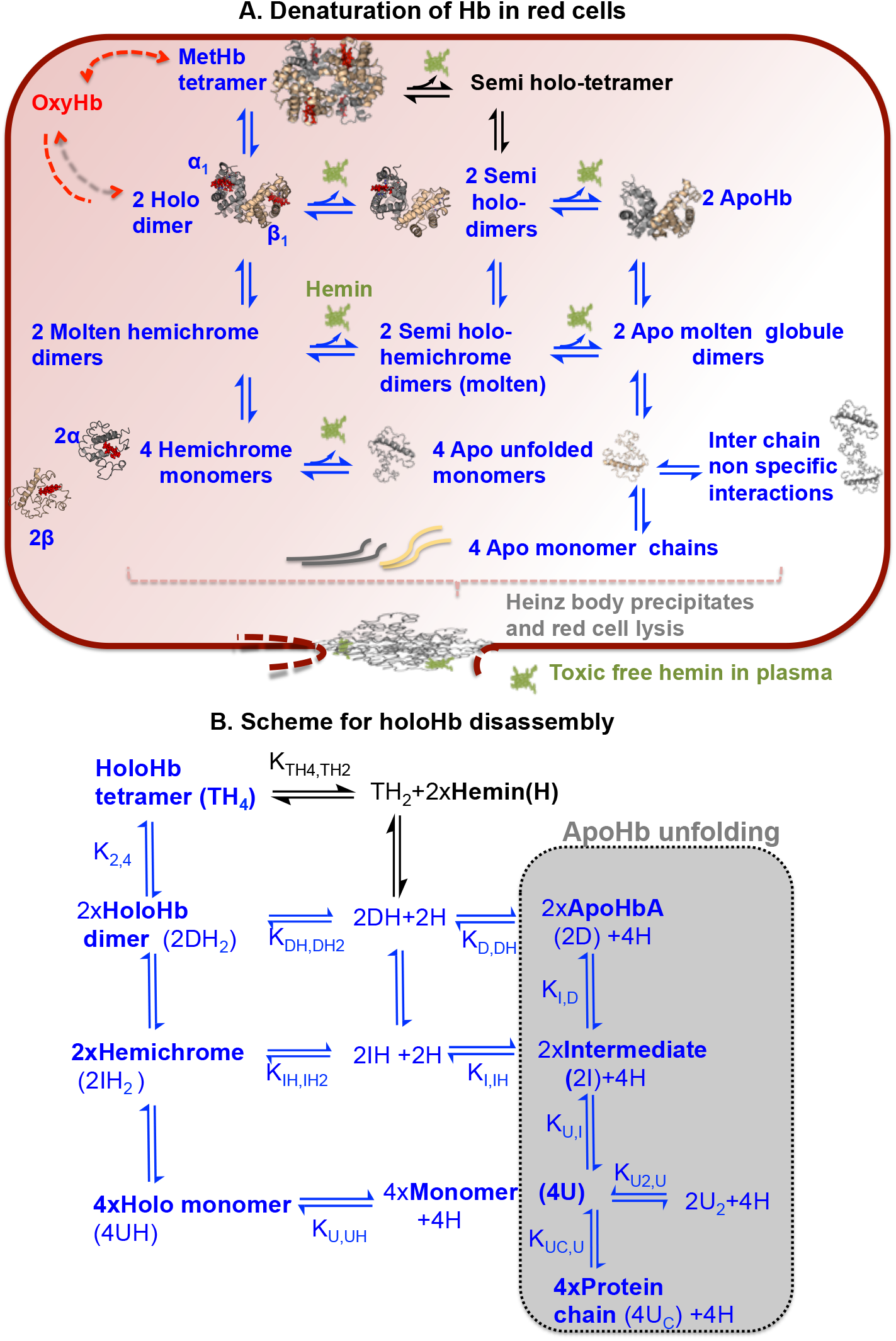
Equilibrium disassembly models of HbA. ***A***, structural description of processes occurring in red cells**; *B***, reaction scheme for processes occurring in our *in vitro u*nfolding studies. Model 1 includes all the reactions in blue in panels A and B, whereas model 2 includes the black reactions involving hemin loss from tetramers as well as the blue reactions in panels A and B. In panel A, free hemin is green, and bound hemin is red. Initially, the holoHb tetramer (TH_4_) disassociates into holoHb dimers (DH_2_). Alternatively, TH_4_ can undergo hemin (H) disassociation from the subunits with lower hemin affinity to form the semi holotetramers (TH_2_). DH_2_ can also undergo H disassociation sequentially first to form semi holo-dimers (DH), followed by further H disassociation, leading to apoHb (D) formation. DH_2_ unfolds into holo-molten globules (IH_2_) with hemichrome characteristics. IH_2_ also can lose H sequentially, first to form semi holo-molten hemichromes, followed by formation of the apo-molten intermediates (I). IH_2_ next disassociates and simultaneously unfolds into holomonomers (UH). Loss of H from UH leads to apo-monomers (U) with residual helical content. U can either associate non-specifically with itself to form transient unfolded dimers (U_2_) or completely unfold into polypeptide chains (U_C_). The apoHb unfolding mechanism was derived and verified independently in our previous study(4).

Building on our earlier work (4), we have been able to establish a global model for human hemoglobin A (HbA) disassembly that involves reversible hemin and subunit disassociations at different disassembly stages (Fig. 1). The concept that multiple assembly/disassembly pathways can occur for protein complexes formed by coupling between two protein subunits or between a protein subunit and a cofactor has been explored for other protein systems but not for hemoglobins (5–8). HbA disassembly is characterized by both types of couplings, and our model predicts that the exact pathway will differ markedly between intracellular red cell and acellular blood plasma environments as a result of marked differences in total protein (and heme) concentrations. In order to develop this model, we measured GdnHCl (guanidine hydrochloride) induced disassembly of holoHb at a wide range of protein concentrations. GdnHCl was chosen as a denaturant to avoid irreversible precipitation of hemin (9–11). We then developed a multilayered analysis approach for Hb disassembly based on our recently published measurements and mechanism for GdnHCl induced unfolding of apoHb (4) and the new holoHb unfolding data generated in this study (Fig. 1). Our model was used to analyze simultaneously circular dichroism (CD) measurements of secondary structural changes in both apo- and holoproteins and populations of hemin-containing unfolding species. The later populations were deconvoluted from visible absorbance measurements at different stages of Hb disassembly. Equilibrium disassembly parameters extrapolated to 0 M GdnHCl concentrations were obtained from fitting of our model to measured CD spectral changes and populations of heme-containing intermediates.

Fatal complications in patients with disassembled or misassembled hemoglobins can result from hypoxia, hemolytic anemias, and iron toxicities due to free hemin release into the blood stream (12–19). The pathogeneses of genetic Hb misassembly and instability diseases often originate from accelerated rates of heme loss, disruption of the inter-subunit interfaces, and accumulation of unstable intermediates states occurring either on- or off-disassembly pathways (Fig. 1) (12, 17, 20). The quantitative hemoglobin disassembly mechanism derived in this work provides the framework to examine in detail various adverse events triggered by hemoglobinopathies and to develop targeted therapies.

The heme iron is axially coordinated to the Nε2 atom of the proximal histidine at the 8^th^ position on the F helix. In the reduced state, this pentacoordinate iron can bind oxygen (O_2_), carbon monoxide (CO), or nitric oxide (NO) at the sixth coordination position on the distal side of the porphyrin ring (21). Auto-oxidation of ferrous heme in oxyhemoglobin leads to aqua-metHb, with H_2_0 bound to Fe(III) protoporphyrin IX (hemin). This metHb form has ≥10,000-fold higher rates of heme disassociation than the corresponding reduced Hb complexes (22–24). Thus, the first step in disassembly of hemoglobins *in vivo* likely involves auto-oxidation to the metHb form, which is then followed by hemin dissociation and protein unfolding. Free heme itself is unstable in the presence of oxygen and auto-oxidizes in milliseconds(21, 25). Therefore, during assembly it is the oxidized form of heme that probably binds to nascent apo polypeptide chains before reduction occurs. In our present study we have focused on the mechanisms and intermediates that occur during metHb disassembly.

In their disassembly study of sperm whale holo myoglobin (Mb), which is structurally similar to each Hb subunit, Culbertson and Olson (11) expanded on the apoglobin unfolding scheme originally developed by Baldwin, Wright, Dyson, and their coworkers (26, 27) to include steps involving hemin dissociation from holoprotein species. Our overall goal was to apply Culbertson and Olson’s approach for examining quantitatively the effects of hemin binding on the unfolding and disassembly of human holoHb, which is a much more complicated oligomeric system with two types of subunits. We further applied deconvolution methods on visible absorbance spectra to compute the populations of native metHb, partially unfolded hemichrome intermediates, and free dissociated hemin in varying mixtures of GdnHCl and holoHb samples. Hemichromes or hemochromes (ferrous form) are not native conformers in folded Hb (28). Ferric hemichromes form via bis-axial coordination of hemin iron to amino acids through combinations of Nε atoms from histidines, S atoms from either methionine or cysteine, and phenoxy O atoms from tyrosine (28).

Specific roles and biophysical characterization of the hemichromes that form during different stages of Hb disassembly have not been well characterized (29–32). Our work shows, for the first time, that (a) heterodimeric molten globule hemichromes play both a critical role for α_1_β_1_ dimer interface formation and dampening of hemin disassociation during the early stages of disassembly; and (b) monomeric and dimeric molten globule hemichromes mediate the major pathway for Hb disassembly at high Hb concentrations in erythrocytes. Our prediction of a predominantly hemichrome mediated HbA disassembly pathway in erythrocytes helps to explain why hemoglobinopathies have been linked to accumulation of denatured hemichrome precipitates that aggregate into inclusion or Heinz bodies (Fig. 1A) (12, 17, 20).

Our small angle X-ray scattering (SAXS) studies confirm that heme binding enables hemoglobin to adopt its functionally compact tetramer conformation. Native human apoHb exists only as a dimer following chemical extraction of hemin (4, 33). SAXS intensity measurements were done on both apo- and holo forms of the recombinant crosslinked HbA or rHb0.1. The linker causes the aporHb0.1 to remain as a tetramer (4). However, our SAXS measurements demonstrate that in the absence of heme, the integrity of tetramer interface in rHb0.1 is still disrupted.

Finally, the shift between apo-versus holo-intermediate mediated Hb disassembly can be modulated by mutations at the inter-subunit interfaces. To verify this idea, we examined fetal hemoglobin (HbF) because of its intrinsically greater resistance to denaturation relative to HbA (34). In HbF, γ subunits are expressed in place of the β subunits (1, 35), with amino acid residue differences at the dimer but not the tetramer interfaces. The populations of molten hemichrome intermediates during the disassembly of holoHbF are significantly greater than those of holoHbA, and are explained by the increased resistance of the α_1_γ_1_ dimer interface to disassociation.

## Material and methods

### Cloning, expression and purification of Hb variants

The molecular biology and protein purification workflow for preparation of holoHb samples were described in our previous paper on apoHb unfolding (see (4) and references therein). Native HbA was extracted from expired blood units (Gulf Coast Regional Blood Bank, Houston, TX). Recombinant native HbA and HbF were co-expressed with *E. coli* Met aminopeptidase from pHE2 and pHE9 E. coli expression plasmids, respectively, in JM109 *E. coli* cells (Promega) as described by Shen et al. (36, 37). The plasmids were gifts from Dr. Chien Ho’s research group (Carnegie Mellon University). The aminopeptidase cleaves post-translationally the initiator Met residue preceding the V1 residue in each Hb subunit. Recombinant HbA variants with αH58L/V62F and βH63L/V67F mutations were then constructed through site directed mutagenesis using Agilent QuckChange® Lightning Kits.

rHb0.1 has a single-residue glycine linker covalently connecting the N-terminal of the first α_1_ subunit to the C-terminal of the second α_2_ subunit. The corresponding non-crosslinked recombinant HbA is known as rHb0.0 (38–40). Both rHb0.0 and rHb0.1 have a V1M mutation in each subunit to initiate protein expression (4). rH0.0 and rhHb0.1 were expressed from pDL111-13e and pSGE1.1-E4 expression plasmids respectively in SGE1661 E. coli cells (38–40). These cells and both plasmids were gifts from Somatogen Inc. (later Baxter Hemoglobin Therapeutics).

Protein expression was induced by 0.2 mM isopropyl β-D-1-thiogalactopyranoside in cells either grown in 500 ml terrific broth in flasks for medium-scale recombinant Hb production following protocols developed by Ho’s group (36, 37) or in a Biostat C 20 bioreactor for large scale expression followed protocols developed initially by Somatogen, Inc. (39) and then modified by our group (41). Isolation and purification of native HbA and the recombinant hemoglobins were done as described previously (36, 37, 39, 41). Cell lysates containing Hb were initially purified using a Zn^2+^ charged chelating Sepharose Fast Flow resin column (GE healthcare) because Hb surfaces are rich with His residues that can bind Zn^2+^ (also see (42)). Further Hb purification was done using a Q Sepharose anion exchange column (GE healthcare), and a final purification step was done for the Hbs expressed from the pHE2 and pHE9 plasmids using a S Sepharose cation exchange column (GE healthcare) to eliminate Hbs with uncleaved initiator Met residues. Quality and identify of the purified Hbs were confirmed by: visible absorbance spectra of holoHb forms; electrospray ionization time-of-flight mass spectrometry; and reverse phase high performance liquid chromatography (Agilent 1220 Infinity LC system) analyses (40, 43). All hemoglobin samples were purified and stored in their reduced CO forms to prevent auto-oxidation by flushing cell lysates, purified proteins, and all the buffers with pure CO for at least 30 minutes (44, 45).

### Preparation of metHb, apoHb, and hemichrome standards

MetHb was prepared by oxidizing HbO_2_ samples with a slight excess of potassium ferricyanide. HbO_2_ was initially prepared by flushing HbCO samples with pure O_2_ for 30 minutes in a chilled rotating flask that was exposed to an intense industrial light to promote photodissociation of the bound CO (21, 44, 46).

ApoHb was prepared from metHb according to the method developed by Antonini’s group (47). First, the pH of a concentrated metHb sample was quickly reduced to ~2.2 followed by extraction of disassociated hemin into cold 2-butanone. Apoglobin recovered from the extraction was then buffer exchanged into 10 mM potassium phosphate, 1mM dithiothreitol (DTT) at pH 7. All ApoHb and metHb samples were prepared and examined at ~ 4 °C (4).

Hemichrome standards were prepared in order to obtain their reference spectra (Fig. 2B). Excess imidazole was added to 12 µM HbA to allow for the binding of imidazole to the hemin iron atom, thereby mimicking the structure of a *bis*-histidine hemichrome. The visible absorbance spectra obtained from this species overlapped with the hemichrome spectra generated by the addition of 600 µM sodium dodecyl sulfate to 12 µM HbA (11, 28, 48, 49).

**Figure 2.**
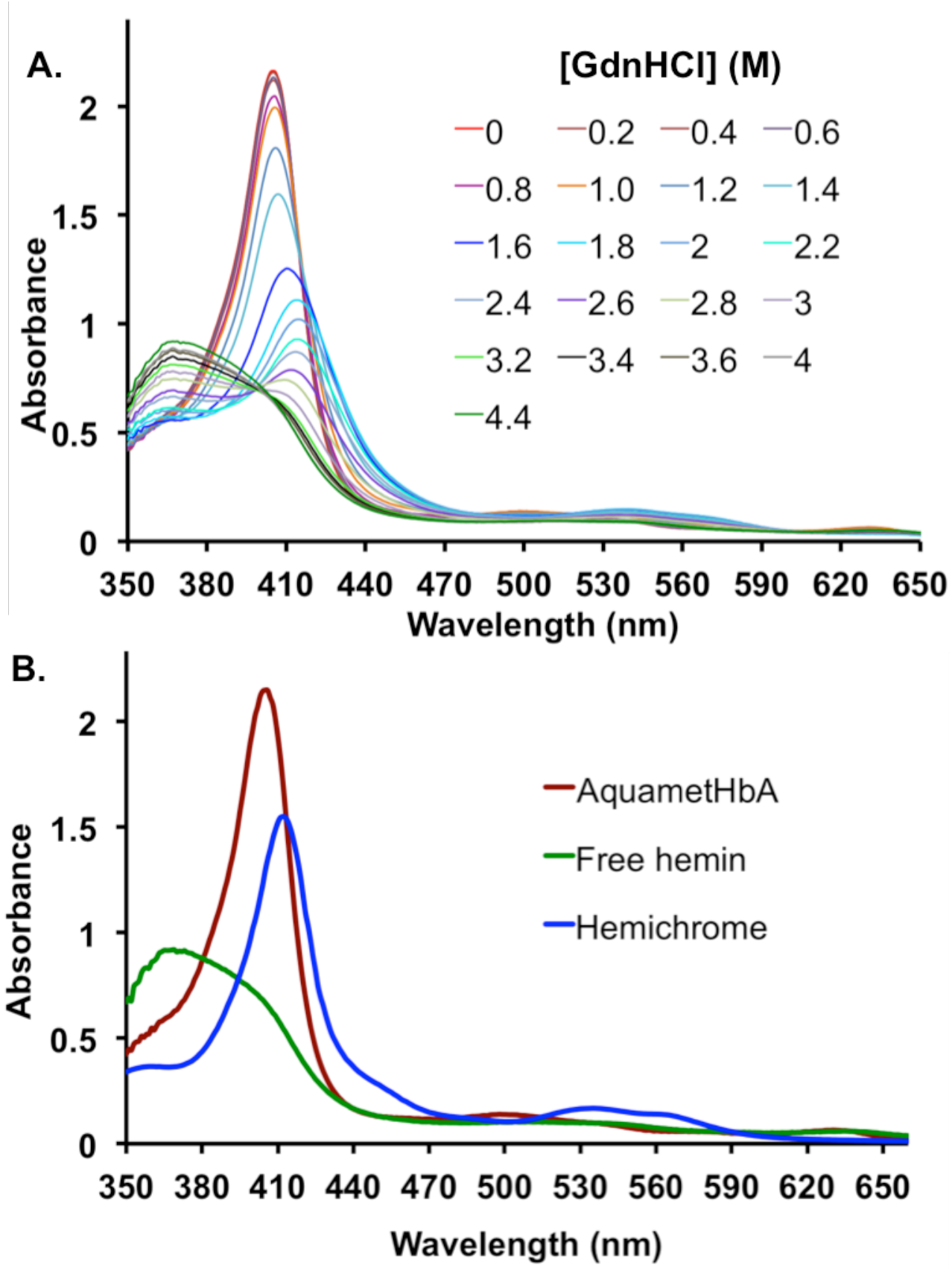
Visible absorbance spectra measurements for HbA disassembly. ***A***, Spectral changes for aquametHbA disassembly induced by GdnHCl; and ***B***, Standard spectra for aquametHbA, hemichrome, and free hemin at 12 µM per subunit protein concentration. Unfolding measurements in panel 2A were done at 12 µM (per subunit) total protein concentration. All holoHb unfolding measurements were done in 200 mM potassium phosphate (pH 7) at 10 °C. Protein samples were initially prepared in aqua-met oxidation states.

### Measurements of holoHb unfolding

MetHb samples were prepared in varying concentrations of GdnHCl in 200 mM potassium phosphate, pH 7. The samples were incubated for 1 hour in a water bath at 10 °C. Spectroscopic measurements of unfolding were then recorded at 10 °C in order to maintain the solubility of unstable folding intermediates. These experimental protocols closely follow our previous apoHb equilibrium unfolding measurements, except that for the addition of 1mM DTT. DTT was added to the apoglobin samples to prevent formation of non-native, disulfide bonds which occur during the preparation and long-term storage of apoHb (4). In our work with metHb samples, DTT was observed to cause partial reduction, large visible absorbance changes, and a complicated mixture of oxidation states, including deoxyHb and HbO_2_. Thus, DTT could not be added to samples for holo-metHb unfolding experiments.

The circular dichroism (CD) and hemin absorbance changes were reversible when holoHb samples incubated in concentrated GdnHCl solutions were diluted to reduce the extent of disassembly and unfolding. CD spectral changes were measured using a Jasco J-810 CD spectropolarimeter, and visible absorbance measurements were measured using a Cary 100 Bio UV-Visible spectrophotometer (4, 44). During the CD spectra data collection, the scanning speed was set to 100 nm/min, data pitch was set to 0.1 nm, data integration time was set to 1 sec, and spectral bandwidth was set to 1 nm. For the visible absorbance measurements, the data collection interval was 1 nm. Spectra for blank buffer solutions were measured initially to enable baseline corrections for both absorbance and CD measurements using instrument software. For a small portion of the visible absorbance measurements, baseline drifts were observed. For each of the spectra with baseline drift, a new baseline was linearly interpolated from the first 21 points of each spectrum at the longest wavelengths using MATLAB R2017a software (The MathWorks, Inc., Natick, MA). The new baseline was then subtracted from the corresponding spectrum.

Visible absorbance spectra at each [GdnHCl] were analyzed in terms of the sum of the matrix multiplication between each species (i) population fraction (D_i_) and the standard basis spectra (B_i_) for the specific i species (Fig. 2B). Three different i species were considered as follows: folded metHb, hemichrome, and free hemin. Visible absorbance deconvolution was carried out by using the fmincon minimizer in MATLAB R2017a (The MathWorks, Inc., Natick, MA) to minimize squared residual difference between the absorbance model spectrum (350nm to 660 nm, 1 nm intervals) at a specific GdnHCl concentration defined in Eq. 1 and the corresponding measured experimental visible absorbance spectrum in order to determine the population fraction (D_i_) of each i species.

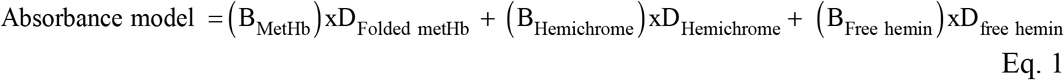

### SAXS sample preparations, measurements, and data analysis

Analytical gel filtration was used to verify whether samples were mono-disperse. Holo- and apoHb samples were prepared by thorough buffer exchange into degassed 10 mM potassium phosphate, pH 7, 5 mM DTT. The buffer was degassed and contained DTT to prevent during data collection error causing bubble formation and radiation damage (RAD) of protein. HoloHb samples were also kept in the CO bound form to further inhibit heme autoxidation during data collection but no DTT was present (44, 50, 51).

SAXS measurements of the Hb samples were done using the Rigaku Bio-SAXS 1000 instrument at the Sealy Center for Structural Biology and Molecular Biophysics at the University of Texas Medical Branch at Galveston, TX. Data were collected at 10 °C, with 12 to 16 hours of exposure to X-ray radiation, monitored hourly for RAD. SAXS measurements of buffer alone were also done to enable blank subtraction. Data collection was done at varying protein concentrations, ranging from 1mg/ml to 6 mg/ml. SAXSLab software (Rigaku) was used for data collection, the SAXNS webserver (http://xray.utmb.edu/saxns) was used for buffer subtraction, and the ATSAS software suite was used for data analysis. From within the ATSAS software, PRIMUS, almerge, and CRYSOL packages were used for preparation and analysis of the observed X-ray scattering curves and GNOM was used to generate pair distance distribution function P(R) plots (44, 52).

## Results

### Disassembly mechanism of holoHbA

In our model for Hb disassembly, heterotetrameric holoHb TH_4_ initially disassociates into two holoHb dimers (DH_2_) without any loss of helical content. In addition to undergoing hemin loss leading to the apoHb dimer (D) and its subsequent unfolding (right sides of Figs. 1A and 1B), the holodimer (DH_2_) can also unfold into a dimeric holo molten globule intermediate (IH_2_), with a hemichrome absorbance spectrum. Each IH_2_ can further unfold and disassociate into 2 holo monomers (UH). We obtained much better fitting to our spectral measurements when we assigned hemichrome absorbance to the UH species. This assignment is supported by previous experimental studies, which showed that isolated aquamet α and β monomers form unstable hemichromes (31, 53). Hemin disassociation from UH leads to formation of apo monomer U states, with residual helical content, and then the completely unfolded chains at very high denaturant concentrations.

Complete chemical extraction of hemin from native HbA leads to a heterodimeric apoHb (D) that has undergone 30% helical content loss (4, 33). Our previous studies show that this D state initially unfolds into a molten globule apo dimer (I), which has lost 70% of its original α-helical content relative to holoHbA. Next, this molten globule apo I state disassociates into unfolded monomers (U) with only residual helical content (≤ 10 %). These monomers then can either interact non-specifically to form transient dimers (U_2_), or completely unfold into polypeptide chains (U_C_) (4). The TH_4_ disassembly scheme in Fig. 1 expands the apoHb model (Fig. 1B, right side, gray section) to include the binding of hemin (H) to the various D, I, and U states and the formation of hemin containing tetramers (TH_4_ and TH_2_ states in Fig. 1). We did not include the formation of tetrameric apohemoglobins in our models because past studies have shown that native human apoHbs can only exist as dimers, and not as tetramers (4, 54–56). The accompanying loss of α-helical content in apoHb due to hemin dissociation is predicted to be occurring around the heme cavity region that also forms the tetramer (α_1_β_2_) interface and thereby causes Hb disassembly into dimers (4, 33).

Previous studies have shown that native metHbA undergoes non-cooperative hemin disassociation, with β subunits having a faster hemin loss rate and presumably lower hemin affinity (22, 57–59). Therefore, we also incorporated non-cooperative hemin binding into our model (Fig. 1) by taking into account sequential hemin disassociation from the DH_2_ and IH_2_ states. However, we made no assumptions about the relative hemin affinities of the two different types of subunits and assigned an equilibrium constant for each sequential dissociation step. In our model fitting, differences between the subunits would be manifested in the extent of the differences between the first and second hemin dissociation or association equilibrium constants.

In our model, hemin (H) disassociates first from one subunit in DH_2_, resulting in dimeric semi-holoHb (DH) formation, and then hemin disassociates from DH leading to the apoHb D state. Similarly for IH_2_, hemin disassociation occurs first from one subunit leading to initial formation of a semi-hemichrome (IH), and then the next dissociation leads to formation of the apo molten globule (I). Past studies have suggested the semi-holohemoglobin formation occurs by either hemin disassociation from the β subunits in holo tetramers/holo dimers or alternatively by initial hemin binding to α subunits in apoHbs (58, 60).

Given the weak affinity of hemin for unfolded proteins (9, 11, 61, 62), the lack of structural evidence about unfolded states, and the limitations of spectroscopic measurements, we assumed no differences in hemin affinity between α versus β variants of U monomers (K_U,UH_) (Fig. 1). Non-specific hemin binding to the U_2_ and U_C_ could not be accurately defined because only free hemin visible absorbance spectra could be observed after the loss of all helical content in the unfolded monomers.

All of these reversible disassembly stages were incorporated into our overall scheme and are represented by Model 1 (blue arrows) in Figs. 1A and 1B. We also considered an additional step involving semi-holo tetramer formation, which was incorporated into Model 2 (both blue and black arrows in Figs. 1A and 1B). In this second model, TH_4_ undergoes disassociation of two hemin molecules initially from the subunit type with lower hemin affinity to form tetrameric semi-holoHb (TH_2_), which then dissociates into a semi-holo dimer (DH).

### Spectroscopic measurements of holoHb disassembly

GdnHCl-induced disassembly of metHbA (Figs. 2 and 3) and metHbF (Fig. 4) was examined as a function of total protein concentration by measuring simultaneously both heme visible absorbance and protein CD spectral changes. With increasing denaturant concentration, holoHb unfolds and undergoes both hemin disassociation (Figs. 2, 3C, and 4C) and loss of α-helical content (Figs. 3D and 4D). CD signal changes at 222 nm were used as a measure of the fraction of α-helical content present in the protein relative to that of folded holoHb (Figs. 3D and 4D (4)). The changes in hemin absorbance spectra were used to compute populations of the various holo species during disassembly and free hemin (H).

**Figure 3.**
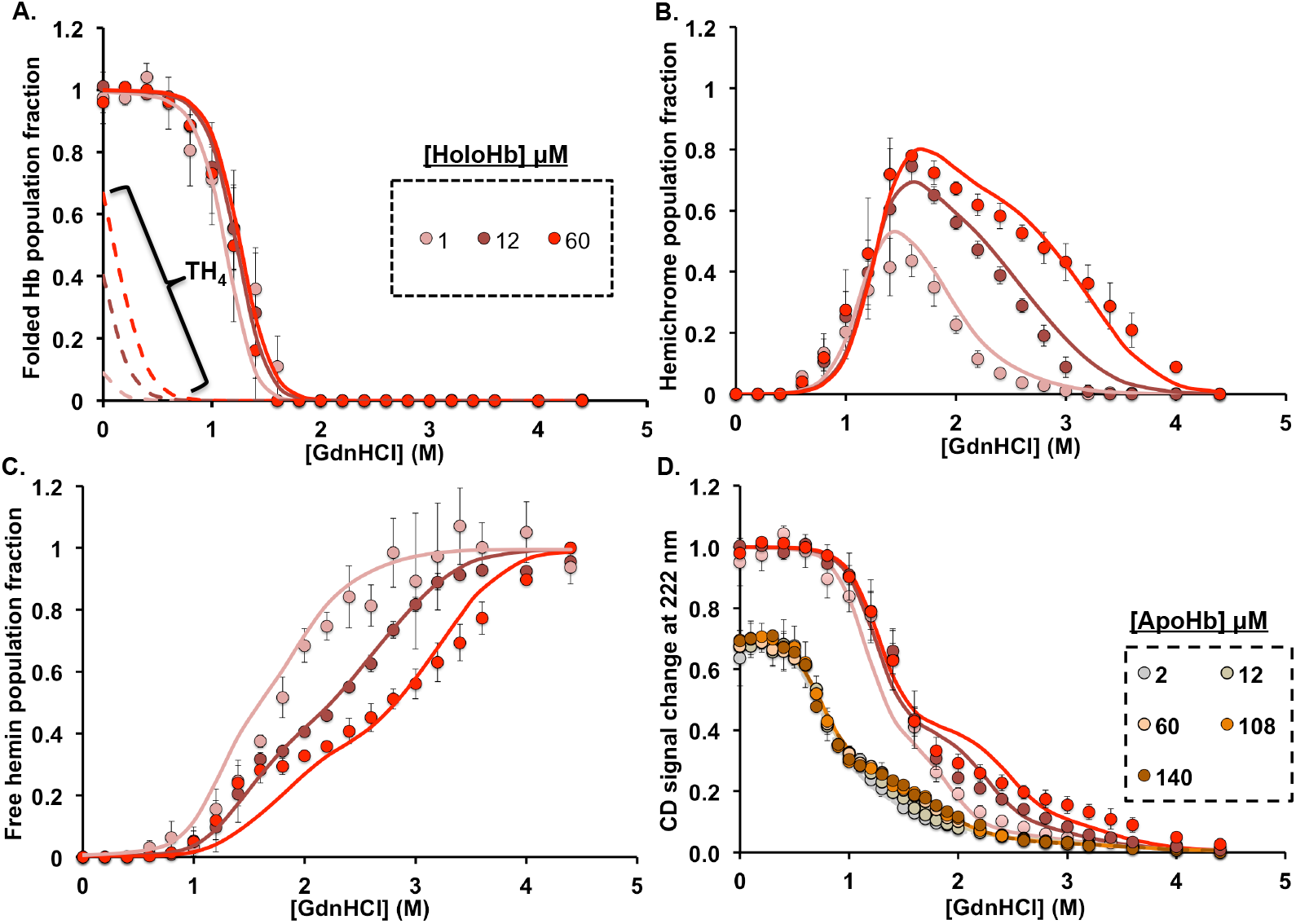
Fitting of holoHbA GdnHCl-induced disassembly measurements to equilibrium disassembly model 1. The solid circles are the measured data at different per subunit total protein concentration and the solid lines are the fits to model 1. The dashed lines in panel A are the theoretical population fraction of folded holoHbA tetramers (TH_4_) derived from the fittings. ***A***, Population fractions of aquamet-holoHb dimers and tetramers. ***B***, Population fractions of hemichromes. ***C***, Population fractions of free hemin. ***D***, Fractional CD change at 222 nm, which corresponds to the change of alpha-helical structure content relative to folded holoHbA. The apoHbA unfolding data (curves at the far left) were previously published (4). The experimental measurements in panels A-C were obtained from deconvolution of visible absorbance spectra recorded between 350 nm and 660 nm. All holoHb unfolding measurements were done in 200 mM potassium phosphate (pH 7) at 10 °C. Protein samples were initially prepared in aqua-met oxidation states. Each data point is an average of triplicate titrations of protein with denaturant.

**Figure 4.**
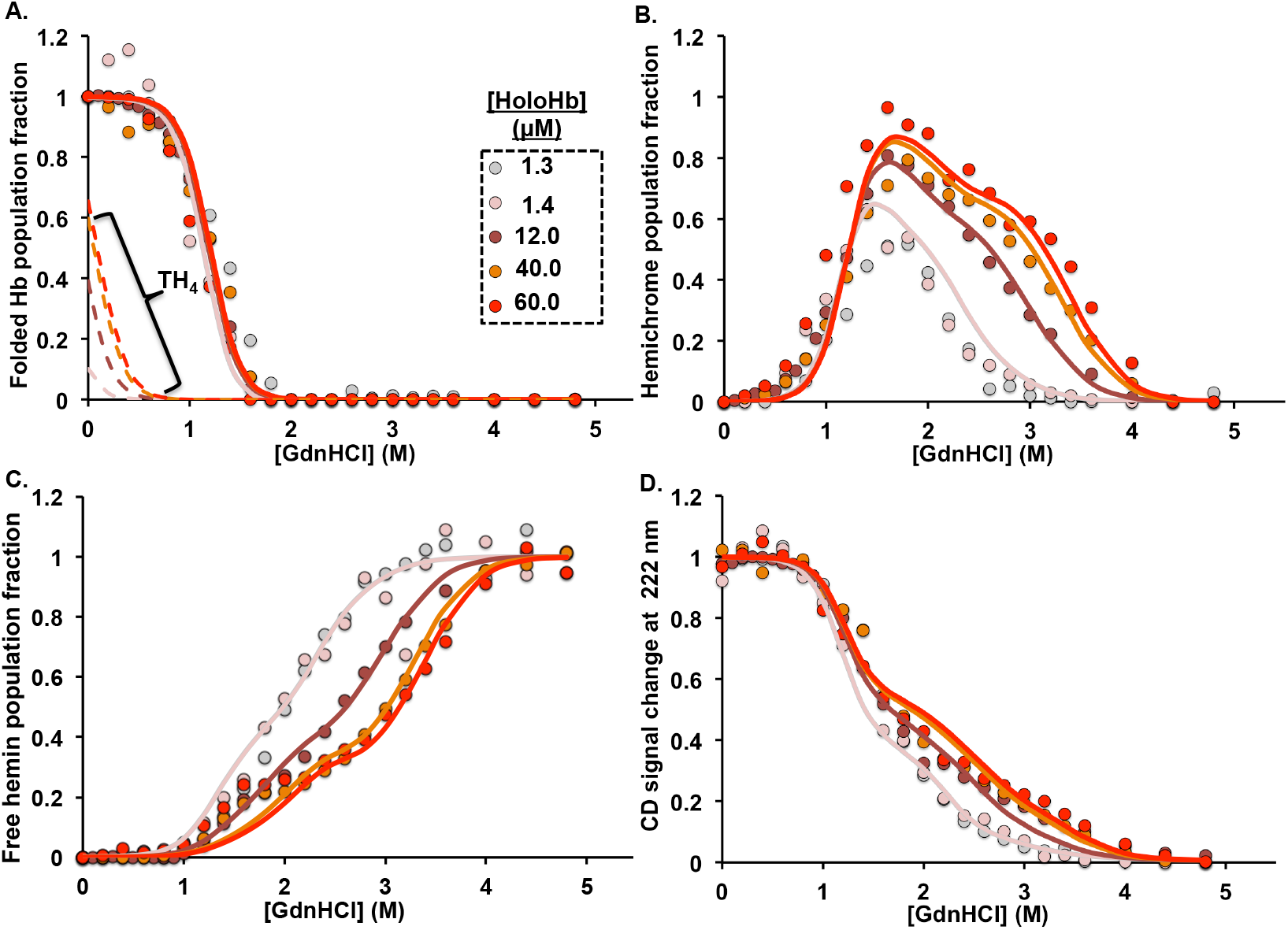
Fitting of holoHbF GdnHCl-induced disassembly measurements to equilibrium disassembly model 1. The solid circles are the measured data at different per subunit total protein concentration, solid lines are the fits, and dashed lines are the theoretical population fraction of folded holoHbF tetramers (TH_4_) derived from the fittings. A. Population fractions of aquamet-holoHb dimers and tetramers. B. Population fractions of hemichromes. C. Population fractions of free hemin. D. Fractional CD change at 222 nm, a measure in the change of alpha-helical structure content relative to folded holoHbF. The experimental measurements in A-C panels were obtained from deconvolution of visible absorbance spectra recorded between 350 nm and 660 nm. All holoHb unfolding measurements were done in 200 mM potassium phosphate (pH 7) at 10 °C. Protein samples were initially prepared in aquamet oxidation states. The 12 µM Hb CD unfolding measurement was obtained from previously published data (4).

As shown in Fig. 2A for metHbA, we observed dramatic shifts in the heme visible absorbance spectra as holoHb disassembles with increasing GdnHCl concentrations. Initially, at low [GdnHCl], the observed visible absorbance spectra were characterized by features that are unique to high-spin aquametHb (Fig. 2B), including a sharp, dominant Soret absorbance peak at 405 nm and additional smaller peaks at 500 nm and 630 nm. At high [GdnHCl] when only residual (≤ 10%) or no alpha-helical content remains (Figs. 2A and 3D), the absorbance spectra were characterized by features that are unique for high-spin free hemin in solution, including a broad Soret absorbance peak around 370 nm and additional smaller peaks around 510 nm and 630 nm (Fig. 2B) (11, 21, 63–65). During intermediate stages of Hb disassembly, another distinct absorbance spectrum appears and is indicative of a low-spin hemichrome (Figs. 2, 3C, and 4C). Hemichrome absorbance spectra are characterized by a distinct and narrow red-shifted Soret absorbance peak around 413 nm and Q bands or visible region peaks at 535 nm and 565 nm (Fig. 2B)(11, 28, 48, 66).

As described in Methods, we deconvoluted the heterogeneous visible absorbance spectral curves at intermediate [GdnHCl] using the three standard spectra for native aquamet, hemichrome intermediates, and free hemin shown in Fig. 2B. The differing population fractions of these species were computed as the GdnHCl titrations proceeded, and the results for aquametHbA are shown in Panels 3A, 3B, and 3C and for aquametHbF in Panels 4A, 4B, and 4C. The hemichrome population reaches a maxima at the beginning of the 2^nd^ phase of unfolding, corresponding to when the protein has lost ~60 % of its alpha-helical content and forms a molten globule (Figs. 2A, 3B, 3D, 4B, and 4D).

### Hemichrome mediated resistance to hemin disassociation and subunit interface disassembly

As shown for HbA in Fig. 3D, the CD curves shifted towards higher [GdnHCl] in the presence of bound hemin in both HbA and HbF (see also (4)). This shift demonstrates the higher stability of holoHb relative to apoHb (4). Previously, we had determined that during apoHb unfolding, the 1^st^ phase involves unfolding of the heme pocket while the 2^nd^ phase corresponds to the dissociation of the heterodimeric molten globule (4). For apoHbs, the protein concentration dependence of the 2^nd^ phase of unfolding demonstrated that it represents disassociation of the α_1_β_1_ and α_1_γ_1_ dimeric molten globules into unfolded monomers (4). The same interpretation applies to the unfolding curves of holo-HbA and HbF in this study. Again, increasing protein concentration shifts the second phase for loss of α-helical content to higher [GdnHCl] (Figs. 3D and 4D). Similarly, the loss of hemichrome populations (Figs. 3B and 4B) and the appearance of free hemin (Figs. 3C and 4C) are shifted towards higher [GdnHCl], demonstrating that there is greater resistance to hemin disassociation with increasing protein concentration. These CD and visible absorbance results demonstrate that bound hemin in the hemichrome state strengthens the α1β1 and the α1γ1 interfaces in the molten globule intermediate.

In our previous studies on apoHb unfolding, we determined that the α_1_γ_1_ dimer interface in HbF is more resistant to dissociation than the α1β1 interface in HbA, with a monomer to dimer association constant (K_UI_) of 8.4×10^9^ M^−1^ for HbF compared to 6.6×10^8^ M^−1^ for HbA (4) (Table 1). Correspondingly, the HbF hemichrome population showed a higher resistance to hemin disassociation during the 2nd phase of unfolding than the HbA IH and IH_2_ populations (Figs. 5B and S1B). rHb0.1, the recombinant variant of HbA with a genetic cross-linker between the α subunits, has a significant hemichrome population at high [GdnHCl], which is greater than that for both HbA and HbF (Fig. 5B). The covalently crosslinked aporHb0.1 tetramer has been previously shown to be much more resistant to disassembly during the second phase of unfolding because two α_1_β_1_ interfaces have to be disrupted in rHb0.1 to generate 2 unfolded **β** monomers and a di-α subunit (4).

**Table 1.**
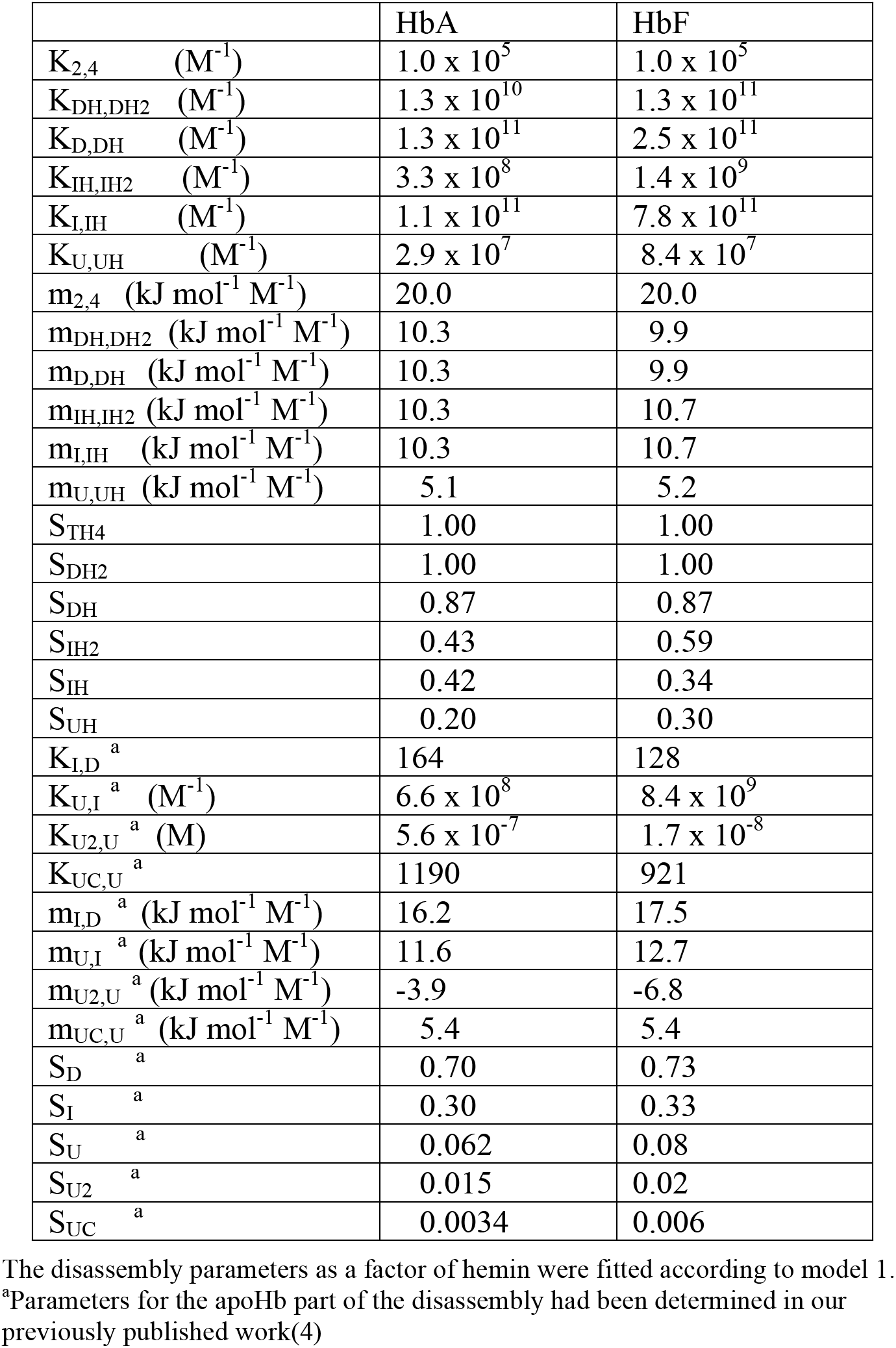
Fitted equilibrium disassembly parameters for HbA and HbF

**Figure 5.**
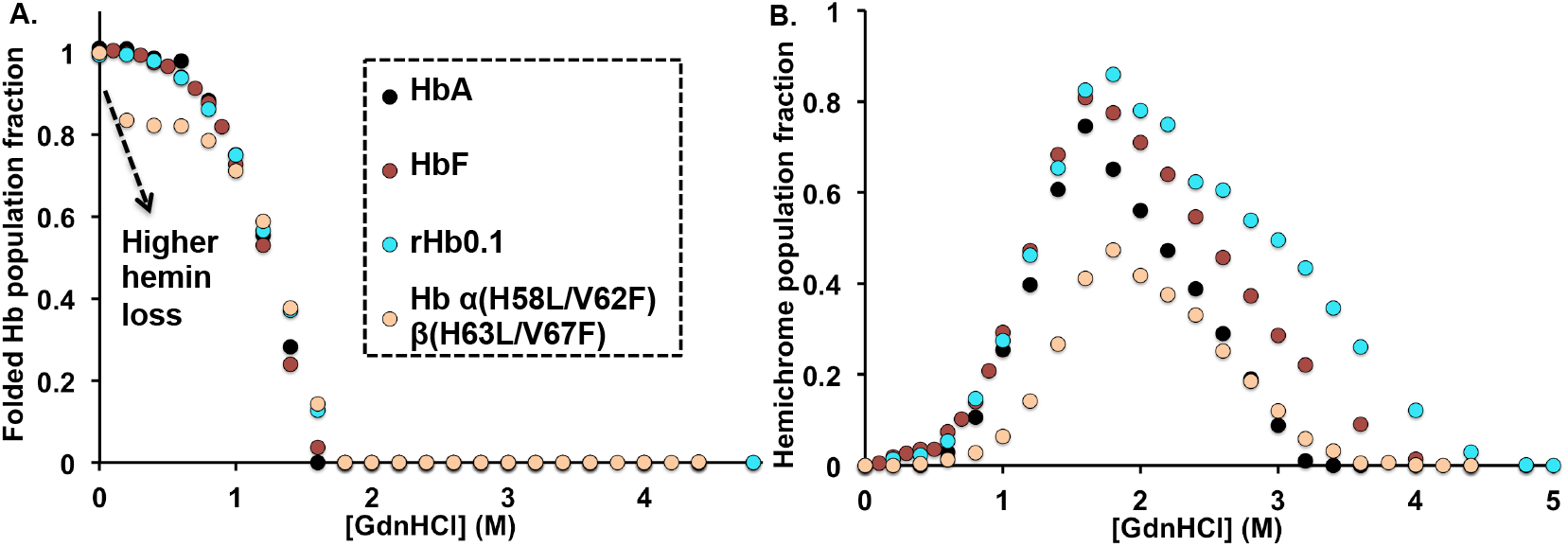
Comparison of GdnHCl-induced equilibrium unfolding measurements of holo-HbA, HbF, rHb0.1, and Hb α(H58L/V62F)β(H63L/V67F). The solid circles are the measured data at 12 µM per subunit total protein concentration. A. Population fractions of met-holoHb dimers and tetramers. B. Population fractions of hemichromes. The experimental measurements were obtained from deconvolution of visible absorbance spectra recorded between 350 nm and 660 nm. All holoHb unfolding measurements were done in 200 mM potassium phosphate (pH 7) at 10 °C. Protein samples were initially prepared in met oxidation states.

For both holo-HbA and HbF, protein concentration dependency was not observed during the 1^st^ phase of unfolding either in the CD curves or the population fractions of the heme spectral species (Figs. 3C, 3D, 4C, and 4D). Hemichromes start to appear during the 1^st^ phase and, in fact, seem to be playing a key role in reducing hemin loss during the early disassembly stages for both HbA and HbF (i.e. little free hemin is present until [GdnHCl] ≥ 1.5 to 2 M). If significant hemin disassociation had been occurring from folded metHb or the newly formed hemichromes, we should have observed population shifts in the curves for native aquametHb and changes in the first phase for the appearance of free hemin as protein concentration increased (Figs. 3A, 3C, 4A, and 4C). Instead, conformational changes around the heme pocket region during the 1^st^ phase promoted a high-spin metHb to low-spin hemichrome transition with heme still bound in the protein. This conformation change is likely induced by the loss of α-helicity around the heme pocket region and is similar to what occurs during the 1^st^ phase of apoHb unfolding (4).

We initially expected to see a dependence on protein concentration for the initial phase of the CD change due to holo tetramer to holo dimer dissociation. However, none was observed and is due to the large tetramer to dimer dissociation constant of aquametHb, which is ~10 µM (67). The threshold of the CD photomultiplier limited the range of holoprotein concentrations to be examined. Within the smaller range that we could examine (1 to ~60 µM total heme), our modeling (Fig. 1) predicted that the tetramer to dimer disassembly was occurring at very low denaturant concentrations before any significant heme pocket unfolding or hemin dissociation occurred. However, if we could have carried out experiments in the millimolar hemoglobin concentration region, our model does predict that we would observe protein concentration effects during the first phases of disassembly (see the results section, “*Modeling HbA disassembly in erythrocytes and in blood plasma*”).

The E7 histidine is the nearest amino acid that can coordinate with the hemin iron at the distal side of the porphyrin ring in Hb. His-E7 often stabilizes exogenous ligands bound to heme iron through hydrogen bonding with the Nε2 atom. However, the Nε2 atom is still at a non-bonding distance of ~4.1 Å from the hemin iron in native metHbA, and a water molecule is coordinated instead (see PDB 3P5Q (68) and (28)). Although we assume the major hemichrome species during native HbA disassembly is most likely due to hemin iron hexacoordination to both the F8 and E7 histidines, to the best of our knowledge, no direct structural evidence for this assumption exists.

Different combinations of axial ligands for hemichrome formation can occur because hemichrome spectra are still observed for the recombinant metHb α(H58L/V62F)β(H63L/V67F) quadruple mutant, in which both distal histidines were replaced with leucines (Fig. 5B). However, the population fraction of the hemichrome intermediate formed during disassembly of the mutant is roughly half that observed for native HbA, implying less stable hexacoordination (Fig. 5B). The standard basis spectrum for folded recombinant metHb α(H58L/V62F)β(H63L/V67F) is different from that of aquametHbA because of the absence of coordinated H_2_O at the hemin iron (69). For the mutant relative to aquametHbA, the Soret peak is blue-shifted to 400 nm and the smaller peak at 630 nm is shifted to 600 nm (Fig. S2). In contrast, the hemichrome spectrum for the LeuE7 containing HbA mutant is very similar to that for the HbA hemichrome. This mutant also showed increased hemin disassociation during the initial phase of disassembly (Figs. 5A and S1B). Past studies of His-E7 to Leu mutations in both Hbs and Mbs attributed the marked increases in rates of hemin dissociation to loss of hydrogen bonding between the His-E7 side chain and the coordinated H_2_O molecule (70, 71). Our new results suggest that the role of hemichrome formation should also be considered when interpreting rates of hemin dissociation for different Hb variants since hemichromes are clearly intermediates in holoHb disassembly.

### Deriving disassembly parameters

The assembly constants defined in Fig. 1B and listed in Table 1 were obtained by extrapolation to 0 M GdnHCl concentration or x, and are represented by the K^0^ values in Eqs. 2 and 3. This extrapolation was done using the dependence of the K constants on [GdnHCl] or x as shown in Eqs. 2 and 3. The apoHb folding constants in Table 1 and eq 2 were taken from our earlier work (4). The m values describe the dependence of the assembly free energies on [GdnHCl] and absolute temperature (T). The K parameters containing H subscripts in Eq. 2 define hemin binding to the various apo subunits in Fig. 1B (except K_TH4,TH2_ which defines hemin disassociation), and K_2,4_ defines the holo metHb dimer to tetramer association equilibrium constant.

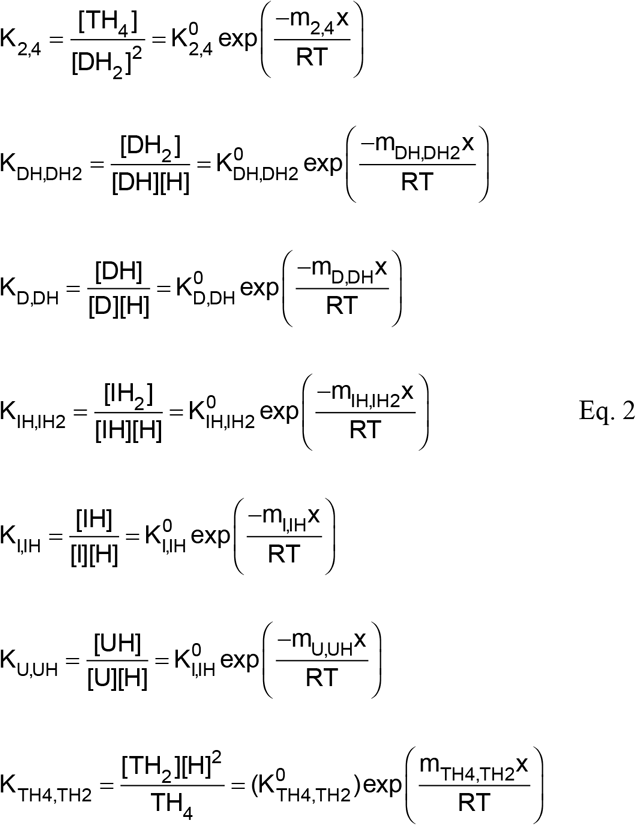

For our overall Hb disassembly/assembly scheme described in model 1 (Fig. 1), the theoretical values of total protein concentration [P] on per subunit basis and total free hemin concentration [Free hemin] in solution were computed from the concentrations of the various species in Eqs. 4 and 5, respectively. This theoretical [P] value should equate to y, the experimentally defined Hb concentration on a per subunit basis. Because there is no excess hemin in the original solutions, [free hemin] in Eq. 5 will equate to [H] in Eq. 2. [Free hemin] in Eq. 5 is computed from the sum of the concentrations of the apo subunits in all the apo- and semi-holo species. For the alternative model 2 that included semi-holo tetramers, [P] and [free hemin] were re-derived with the addition of the terms 4[TH_2_] and 2[TH_2_] to the right sides of Eqs. 4 and 5 respectively.

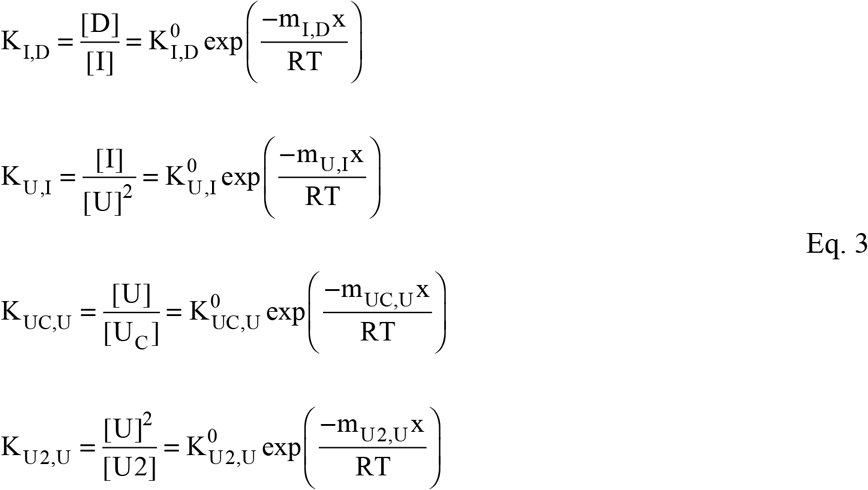

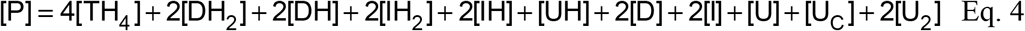

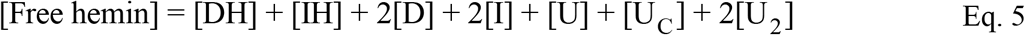

The general population fraction Y_i_ on per subunit basis for a specific folding species (i) is described in Eq. 6, where specific examples are given for the free hemin fraction (Y_H_) and holoHb tetramer population fraction (Y_TH4_) from model 1. The total population fraction of folded holoHbs, encompassing both the holoHb dimers and tetramers, is defined as Y_Folded holoHb_ =Y_TH4_ + Y_DH2_ + 0.5Y_DH_ for model 1, and Y_Folded holoHb_= Y_TH4_ + Y_DH2_ + 0.5(Y_TH2_ + Y_DH2_) for model 2. In both models, the total population fraction of hemichromes was evaluated as Y_Hemichromes_ = Y_IH2_ + 0.5Y_IH_ + Y_UH_.

The theoretical Y_Folded holoHb_ value was assumed to correspond to the population fraction of native metHb spectral species (red spectrum in Fig. 2B) derived experimentally from deconvolution of the measured visible spectra as a function of [GdnHCl] (Eq. 1). Y_Hemichrome_ and Y_H_ were assumed to correspond, respectively, to the fractions of low spin hemichrome spectrum (blue spectrum in Fig. 2B) and high spin free hemin spectrum (green spectrum in Fig. 2B) in the measured visible absorbance data. A weight of 0.5 was assigned to the semi-holoHb Y values in the expressions for Y_Hemichrome_ and Y_Folded holoHb_ to account for the subunits present that already had undergone hemin disassociation. The theoretical normalized total CD signal, S_Total_, represents the fractional α-helicity measured at 222 nm relative to that for folded holoHb. S_Total_ was defined as the sum of the product of Y_i_ and the intrinsic normalized CD signal S_i_ for each folding species (not including free hemin) as shown in Eq. 7. S_i_ is independent of x and y and is defined as the fraction of α-helical content of species i relative to folded holoHb.

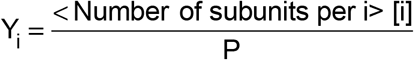

e.g. for Model 1

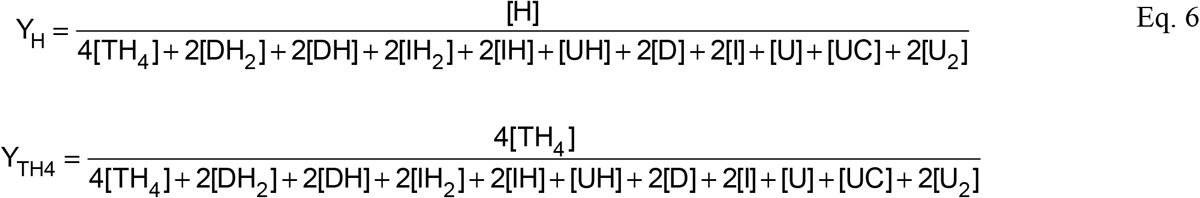

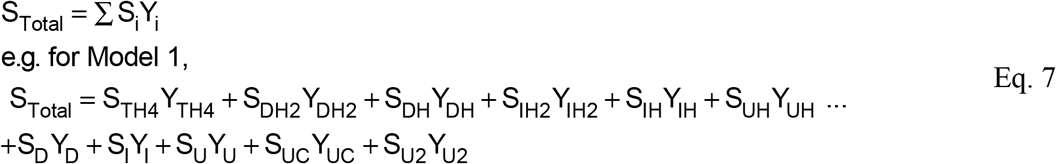

The K, m, and S_i_ values characterizing holoHb unfolding were estimated by global fitting of (a) specific Y_i_ or sums of Y_i_ values to the corresponding measured populations of native metHb, hemichrome, and free hemin obtained from deconvolution of the visible spectral data; and (b) S_Total_ to the experimental CD values that were normalized against the α-helical content of folded holoHb (Figs. 3, 4, and Table 1). The fits to these experimental measurements were done as a function of x (GdnHCl concentration) and y (total protein concentration) variables for both holoHbA and holoHbF. The K, m, and Si values for the apoHb unfolding steps were fixed to the values determined from our previous, independent apoHb unfolding studies (4, 11). The m values for the association constants for hemin binding to folded heme pockets were set equal to each other (e.g. m_DH,DH2_ = m_D,DH_), and the m values for the parameters describing hemin binding to the molten dimers were also set equal to each other (e.g. m_IH,IH2_ = m_I,IH_) (Table 1, Table S1).

The K_2,4_ value for metHb dimer-dimer association was initially set to 1×10^5^ M^−1^ for metHbA, based on previous studies. Because there is no protein sequence difference at the tetramer interface between HbA and HbF, the same K_2,4_ value was used for HbF (67). This number did not change much during the model fittings. The lack of change was likely due to: (a) the absolute value of K_2,4_ being poorly defined but in the 1×10^5^ M^−1^ range; and (b) tetramer dissociation being complete at very low denaturant concentrations ([GdnHCl] ≤ 0.5 M).

The Y_i_ values had to be computed first by defining them in terms of the K constants at different x and y values (left sides of Eqs. 2 and 3) for comparison with the experimental data. For this purpose, [U] and [H] values at a given x and y were obtained by numerically applying Newton’s method (72) for systems of non-linear equations (Eqs. 8–9) expanded from the definitions of [P] (e.g. Eq. 4) and [Free hemin] (e.g. Eq. 5). As shown for Model 1 in Eqs. 8–9, all species were expanded only in terms of [U], [H], and K values. The initial values of [U] and [H] obtained numerically were then used to compute new values of Y_i_.

Starting from these initial estimates of Y_i_, lsqcurvefit function (a non-linear least-squares solver) in MATLAB R2017a (The MathWorks, Inc., Natick, MA) was applied to globally fit the theoretical models to the experimental measurements as shown in Figs. 3, 4, and S3. An algorithm was written in MATLAB R2017a (The MathWorks, Inc., Natick, MA) to apply the numerical analysis method and fitting function in an iterative loop where the K, m, and S_i_ parameters updated from each optimization loop was fed back to the numerical analysis part of the algorithm to obtain a new sets of [U] and [H] values. The implementation of the loop is stopped when the best fit between the experiment and model is reached as defined by a minimum steady value of χ2, the squared residual difference between the models and experimental measurements.

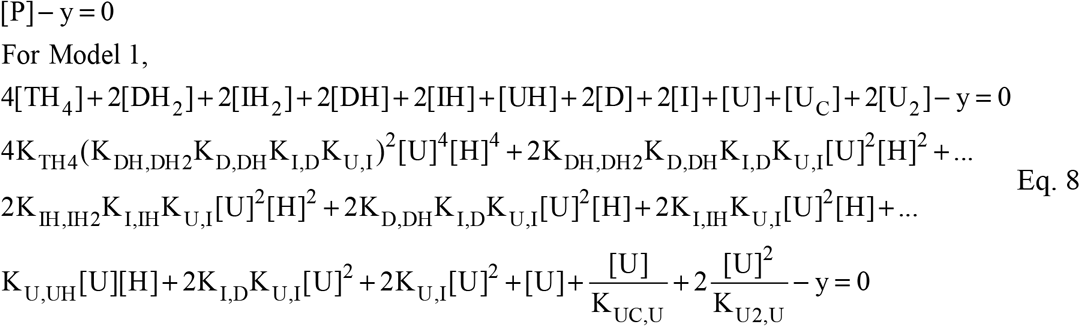

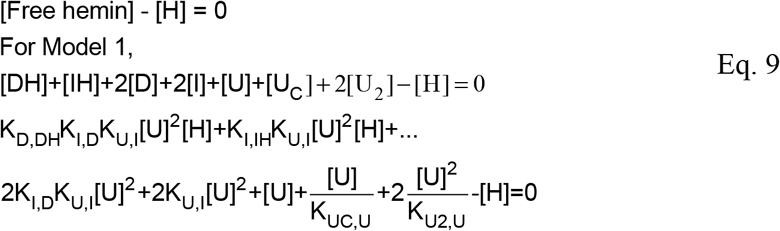

Our overall disassembly Model 1 (Fig. 1, blue arrows) was fitted to HbA unfolding data (Fig. 3) using the optimized parameters in Table 1. A χ2 value of 0.76 was obtained.

Model 2 was proposed in order to try to determine the hemin binding constant for the low affinity subunit (probably β chains) in tetrameric HbA (Fig. 1). When HbA unfolding data were fitted to Model 2 (Fig. S3) with 2 additional degrees of freedom (Table S1), there were only small changes in the goodness of the fit (χ2=0.65). More importantly, the population fraction of semi-holo tetramers, with hemin disassociated from the low affinity subunits, was always close to 0 (Fig. S3A inset). Even this more complex scheme indicated that small amounts of GdnHCl were sufficient to disrupt the weak holo tetramer interface before any significant unfolding or hemin dissociation occurs at the protein concentrations that were accessible to our spectral measurements (dashed lines in Figs. 3A and S3A).

When Model 2 was simulated later at high protein concentrations (~mM range) (see next section and Fig. S4A), the population of semi-holo tetramers was still almost 0. This result also suggests that semi-holo tetramers are not a predominant species during HbA assembly, and heme insertion is occurring much earlier during dimer assembly (see Discussion). Furthermore, other kinetic studies have documented that hemin loss rates increase dramatically when Hb tetramers dissociate into dimers (22, 24, 73). Thus, we chose to consider only the simpler Model 1, in which complete dissociation of holo tetramers into dimers occurs before either hemin loss or unfolding.

This model (Fig. 1) predicts sequential binding of hemin to the apo dimer with association equilibrium constants equal to K_D,DH_ = 1.3 × 10^11^ M^−1^ for the first step and K_DH,DH2_= 1.3 × 10^10^ M^−1^ for the second step. By considering the binding sites to be non-interacting with each other, we interpret these constants in terms of a two-step Adair binding equation (74) to evaluate the intrinsic hemin affinity per subunit. In this case, the intrinsic hemin affinity per subunit for the first step is 1/2K_D,DH_ or 6.5 × 10^10^ M^−1^, and the intrinsic affinity per subunit of the second step is 2K_DH,DH2_ or 2.6 × 10^10^ M^−1^. This ~3 – fold decrease in hemin affinity could be due to either negative cooperativity or differences in hemin binding to the α versus β subunits. The non-cooperative, subunit differences interpretation is the most reasonable because we and others (22, 57–59) have shown that the rate of hemin dissociation from β subunits is higher than that from α subunits in metHbA. Therefore, these intrinsic hemin affinities can be interpreted to indicate that the initial loss of hemin from the native folded dimer comes from the β subunit generating the semi-holo DH state with hemin still in the α subunit.

Hemin binding to the apo molten HbA dimer (I in Fig. 1B) also occurs sequentially, and in this case, the difference in hemin affinity between the first and second steps is even larger, with K_I,IH_ = 1.1 × 10^11^ M^−1^ and K_IH,IH2_= 3.3 × 10^8^ M^−1^ (Table 1). In this case, the intrinsic subunit affinity for the first step is 1/2K_I,IH_ =5.5 × 10^10^ M^−1^ and almost equal to that for the first step in hemin binding to the native D state. In contrast, the intrinsic affinity for the second step is only 2K_IH,IH2_ =6.6 × 10^8^ M^−1^. This large decrease in affinity suggests the subunit differences increase ~30-fold in the molten intermediate relative to the folded state. Based on the known higher hemin loss rates from the β subunit in the folded metHbA (22, 57–59), it appears likely that its affinity for hemin is even lower in the molten or hemichrome I state.

Unfolded apo- and holo monomers at high [GdnHCl] have only residual α-helical content, ranging between ~10 % and 20 % relative to folded HbA subunits (Table 1) (4). Thus, as expected, the fitted hemin affinity constant for these unstructured monomers is small K_U,UH_= 2.9 × 10^7^ M^−1^ (Table 1) and likely dominated by non-specific interactions (9, 11, 61, 62) (Table 1).

As shown in Fig. 4, we were able to analyze successfully holoHbF unfolding data with the same Model 1. In this case, a somewhat higher χ2 value of 1.76 was obtained. In folded HbF dimers, the hemin affinity for the first and second steps in hemin binding were more similar with K_D,DH_= 2.5 × 10^11^ M^−1^ (intrinsic subunit affinity, 1.3 × 10^11^M^−1^) and K_DH,DH2_= 1.3 × 10^11^ M^−1^ (intrinsic subunit affinity, 2.6 × 10^11^M^−1^) indicating non-cooperative binding and little or no subunit differences. However, the affinities of the first and second steps for binding to the HbF molten intermediate are quite different, with K_I,IH_ = 7.8 × 10^11^ M^−1^ (intrinsic subunit affinity, 3.9 × 10^11^) and K_IH,IH2_= 1.4 × 10^9^ M^−1^(intrinsic subunit affinity 2.8 × 10^9^ M^−1^). Again, it seems that the γ subunits have much lower hemin affinity in the molten globule intermediate. The U monomers for HbF also had ~10-fold higher hemin affinity than in HbA, and the holo monomers, UH had slightly higher α-helical content at 30% relative to folded HbF (Table 1). Regardless of the exact interpretation, it is clear that one of the hemichrome containing subunits in the molten dimer states of both HbF and HbA has a relatively high affinity for hemin, which is similar to that in the folded D state.

### Modeling HbA disassembly in erythrocytes and in blood plasma

During maturation of red blood cells, all of the membrane organelles in the cytoplasm, including the nucleus and mitochondria, are eliminated to allow for dense packing of hemoglobin to a concentration of ~20 mM per subunit (75–77). At these high concentrations, native oxyhemoglobin exists almost completely as holo tetramers because the tetramer-dimer disassociation constant (K_4,2_ or 1/K_2,4_) is on the order of 10^−6^ M, and the corresponding value for deoxyhemoglobin is even lower on the order of 10^−12^ M (67, 78, 79).

If oxidative stress occurs in erythrocytes due to either genetic disorders, environmental chemicals, or drugs, denaturation of the resultant metHb can occur in the cell cytoplasm. At these high hemoglobin concentrations, the major disassembly pathway will be dominated by holoprotein intermediates, represented by the left side of Fig. 1. To illustrate this effect quantitatively, we computed the population fractions of all the intermediates for a GdnHCl-unfolding titration at 20 mM total protein, using the parameters in Table 1 (Figs. 6A and B). The initial step involves dissociation of holo tetramers into holo dimers, which is followed by unfolding into molten holo dimers with hemichrome spectra. This IH_2_ species is the dominant intermediate with small amounts of the semi-holo molten globule, IH. The IH_2_ and IH species dissociate predominately into unfolded holo monomers before any significant hemin loss occurs. The high Hb subunit concentration inhibits dissociation of both the dimeric intermediates and bound hemin. Even the loss of weakly bound hemin from unfolded monomers is inhibited until very high [GdnHCl]. When model 2 and the parameters in Table S1 were used to simulate the population fractions at 20 mM Hb, the same conclusion on the dominant role of holoprotein intermediates is reached (Fig. S4A and S4B).

**Figure 6.**
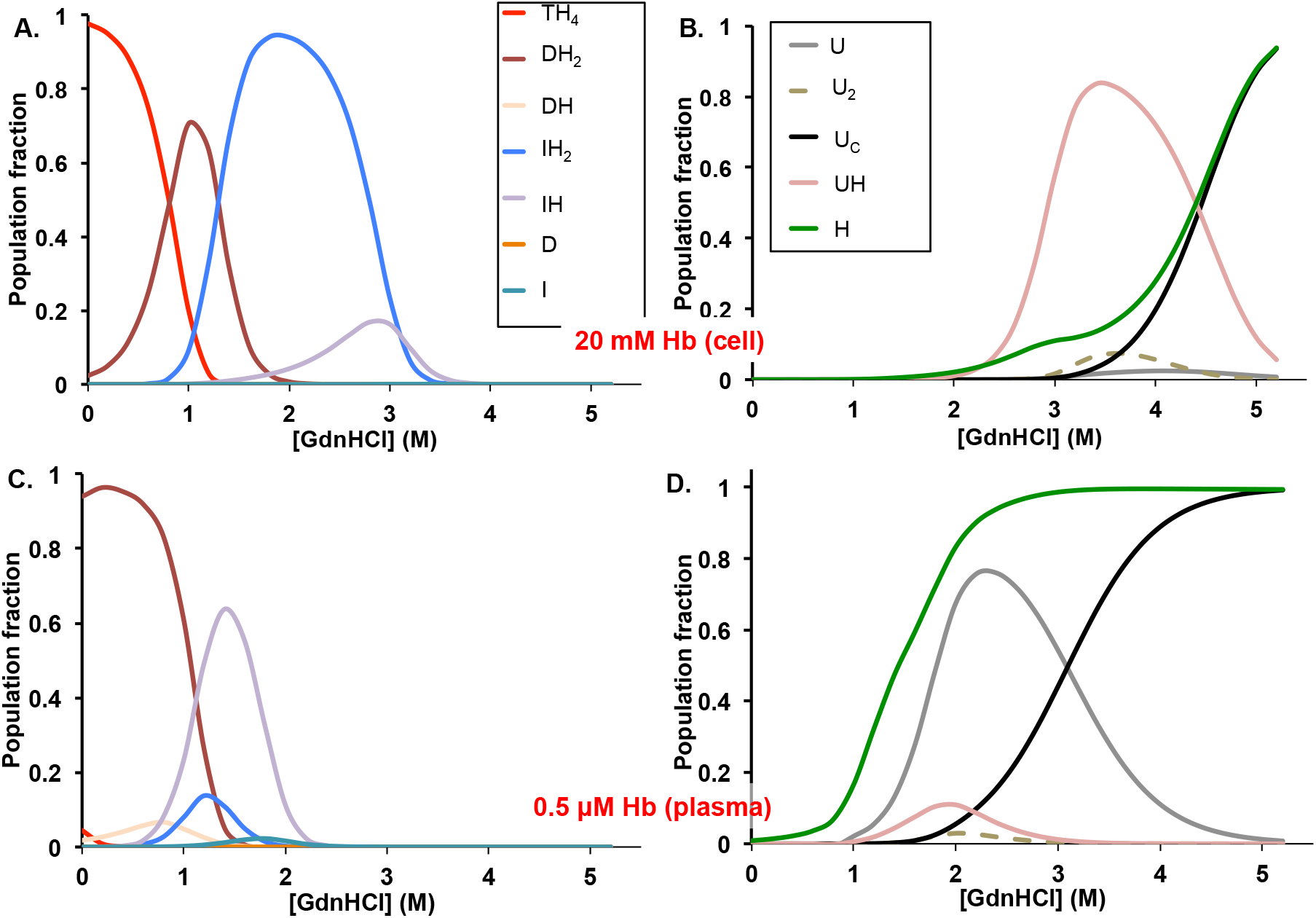
Simulated HbA disassembly population fractions. ***A***, Hb packed red blood cells and ***B***, dilute blood plasma. The results demonstrate Hb concentration directed disassembly pathways. Simulations were done using Model 1 (Fig. 1) and the parameters in Table 1 as a function of [GdnHCl].

Hemolysis of red cells leads to the release of hemoglobin into plasma and can be triggered by hemoglobinopathies, red cell aging, oxidative stress, blood infections by heme iron extracting pathogens, complications during cardiovascular medical procedures, and blood transfusions. Dilution into the large volume of plasma leads to very low, micromolar Hb concentrations, which causes almost complete disassociation of tetrameric Hb into dimers and acceleration of auto-oxidation (23, 80–85). Thus, when hemoglobin is diluted into plasma after red cell lysis, metHb formation occurs and dissociation of both the inter-subunit interfaces and hemin are facilitated. This situation was simulated in Figs. 6C and 6D using the parameters in Table 1 and 0.5 µM total protein. The disassembly pathway is quite different from that in red cells. The initial holoprotein is primarily a dimer, DH_2_ (Fig. 6C) and the dominant intermediates are the semi-holo molten dimer, IH, and the apo unfolded U monomer until the completely unfolded chains, U_C_ are formed. Perhaps the most striking difference between the red cell versus plasma pathways is the early appearance of free hemin before unfolding is finished (Fig 6D). At low Hb concentration, dissociation of the α_1_β_1_ dimer interface is facilitated, resulting in low populations of the molten dimer hemichromes. Hemin dissociation is also facilitated, leading to predominately apo intermediates and unfolded chains even before the end of the simulated titration.

### SAXS analysis of apoHb properties in solution relative to holoHb

SAXS intensity measurements (Fig. 7) were done on both apo- and holo forms of recombinant crosslinked HbA or rHb0.1 and also on the holo form of the corresponding non-crosslinked rHb0.0 (38–40). Crosslinking causes aporHb0.1 to remain a tetramer under conditions where native apoHb is a dimer (4). We chose the aporHb0.1 variant for SAXS analysis in order to have a covalently crosslinked tetramer as standard for the radius of gyration (R_g_) for holoHb tetramers. The SAXS measurements were fitted to the Guinier plot equations (Fig. 7 inset) to obtain R_g_ for each molecule. Minor effects of molecular crowding were only observed for aporHb0.1, occurring at protein concentrations >1.5 mg/ml (Fig. S5). For aporHb0.1, R_g_ and the pair distance distribution function P(r) between atoms was determined with SAXS scattering measurements extrapolated to infinite dilution in order to remove the molecular crowding effects (86) (Figs. 7, S5, and S6). AporHb0.1 at 0.0 mg/ml exhibited a R_g_ of ~29.4 Å, whereas for the holo forms of rHb 0.0 and rHb0.1, the experimental R_g_ values both ~24 Å, as expected from their known crystal structures (e.g. PDB ID 1O1L) (Figs. 7 and S5). Thus, aporHb0.1 is expanded in solution. These kinds of expansions are often observed for partially unfolded proteins (86), and support our view that hemin dissociation causes loss of the ability of hemoglobin to form compact and cooperative tetramers. The losses of helical content are likely occurring around the heme cavity regions, which also forms the α1β1 tetramer interface, allowing expansion of the crosslinked protein.

**Figure 7.**
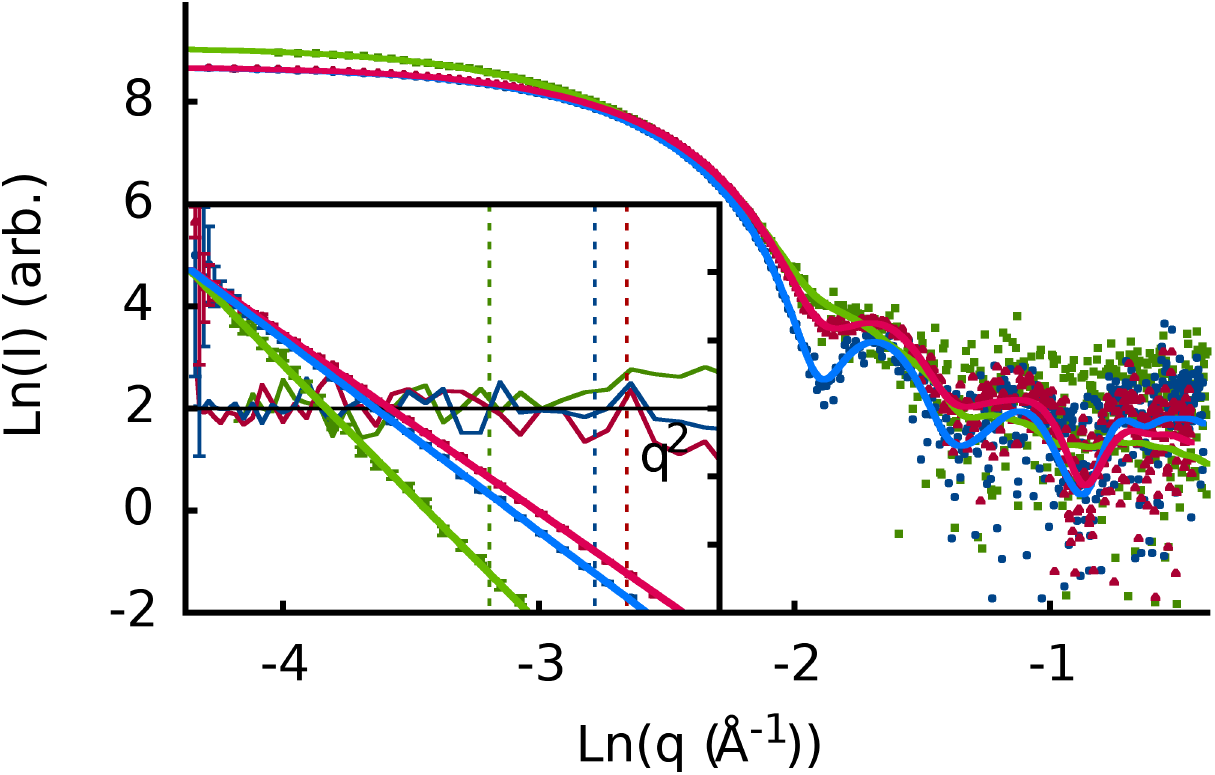
SAXS Log intensity profiles and Guiner plot (inset) for recombinant hemoglobins. For the Log-Log plots, the samples shown are the zero-concentration extrapolated aporHb0.0 (green), 2.0 mg/ml Hb0.1 (blue) and 2.0 mg/ml Hb0.0 (red) Inset: The Guinier plots with their residuals (darker lines) and (1.3/R_g_) Guinier q-upper limits (dashed lines for R_g_=29.4, 24.3, 23.6 Å). The expansion of the apoprotein is obvious from the less distinct minima in the higher angle data.

Interestingly, our SAXS measurements also suggest that this loss of helical content due to hemin dissociation increases the susceptibility of Hb to molecular crowding. Similar effects of molecular crowding probably might occur for the partially disordered assembly intermediates and during early folding events for Hb in vivo. With more sticky regions of unfolded chain segments exposed and expanded in solution, there is also a higher probability they can come in closer contact and even interact weakly with other protein molecules causing the crowding.

## Discussion

### Global disassembly pathway for human hemoglobin

Hetero-tetrameric hemoglobins evolved from monomeric vertebrate myoglobins within the globin metalloprotein family about 600 millions ago (1, 87, 88). In this work, we have shown for the first time that human hemoglobin shares a common disassembly pathway with mammalian myoglobin. For both systems, hologlobin disassembly is mediated via a molten hemichrome intermediate. However, relative to the simpler myoglobin monomer system described (11, 26, 27), the human hemoglobin assembly/disassembly mechanism evolved to be much more complex. This complexity is demonstrated by the multiple steps for heme binding to dimeric intermediates and by the sequential formation of the dimer (α1β1) and then tetramer (α1β2) interfaces (Fig. 1).

To best of our knowledge, no one has previously tried to analyze reversible holoHb disassembly quantitatively with a comprehensive model, probably because of its complexity. In addition, most studies involved thermal denaturation (89–91), which leads to precipitation of both the globin chains and free hemin in conventional buffers. Our study is also the first attempt to measure equilibrium hemin binding parameters for human hemoglobin, and provides reasonable estimates for comparison with previous kinetic studies (22, 90, 92). We were also able to estimate heme binding parameters at different stages of Hb assembly/disassembly, independently, and characterize how they influence inter-subunit assembly and globin folding. Determination of hemin affinity constants is not possible by direct titration of apoglobins with hemin because free hemin stacks to form dimers and higher molecular weight aggregates even in the micromolar region (65). In our experiments, the hemin equilibrium binding constants were obtained by simultaneously analyzing GdnHCl-induced unfolding curves for both apo- and holoHb in order to avoid problems associated with apoglobin and hemin precipitation.

Hargrove, Barrick, and Olson (92) had previously concluded that the association rate constant for hemin binding to apoglobin seems to be relatively invariant and ≈ 1 × 10^8^ M^−1^ s^−1^ per subunit regardless of the globin structure for either apoHb or apoMb. Past kinetic studies were focused mostly on measuring heme disassociation rates from the end-state products of Hb assembly e.g. TH_4_ and DH_2_ (Fig. 1) (22, 93). However, such kinetic studies did not attempt to consider how these rates and hemin affinities change as unfolding proceeds via molten globules with significant reconfigurations of the hemin pockets.

The approaches that led us to successfully resolve “new” holo intermediates and their roles disassembly were: (1) deconvolution of visible absorbance signals into various intermediate hemin containing populations; (2) model fitting to both apo- and holo protein unfolding spectral measurements; and (3) examination of the protein concentration dependency of bimolecular events involving both inter-subunit interactions or heme binding. We also developed a model evaluation approach that incorporates numerical analysis methods to solve for disassembly populations and takes into account the complexity of the Hb protein with four subunits and four bound hemin molecules. For monomeric heme proteins, more straightforward quadratic equations can be used to solve for fractions of various holo- and apo intermediates (11, 63).

### Physiological relevance

The results in Figs. 6A and B can provide insights for the assembly pathway at various protein and heme concentrations during hemoglobin biosynthesis. First, hemin binding is a bimolecular process and required for tetramer assembly. Therefore, at high heme and protein concentrations, the fraction of the holo molten globule and native dimers will outcompete their apo counterparts to promote tetramer formation. In addition, at these high concentrations, weak hemin binding to U monomers might be interpreted as an induced fit mechanism (5), promoting holoprotein assembly from newly expressed α and β chains in immature red blood cells or reticulocytes. This prediction seems to fit with previous observations by Spirin’s group (94), which suggest co-translational heme binding to nascent globins chains coming off the ribosome in cell-free lysates of rabbit reticulocytes. Alternatively, at much lower concentrations of Hb subunits and free heme, hemin binding during assembly will proceed through conformational selection (5) of partially or fully-folded conformations (Figs. 6C, 6D). Even though the total amount of holoHb being generated during erythropoiesis is large, the absolute concentrations of the newly synthesized apo chains and free hemin in the steady state are likely to be low due to rapid Hb assembly and hemin sequestering in membranes and carrier proteins (95). In addition, if unfolded globin chains and free hemin were present at high concentrations, they would self-aggregate rapidly to form protein precipitates and stacked hemin complexes (65), which would compete with assembly, causing unwanted pathological situations.

We do acknowledge that our disassembly model derived from *in vitro* studies cannot completely capture *in vivo* disassembly/assembly mechanisms. Within the packed red-cells, non-specific or quinary interactions (96, 97) will occur between different combinations of α and β chains for formation of transient U_2_ species (Fig. 6B) (4). Our previous *in vitro* experiments have shown that transient aggregation to generate apo U_2_ intermediates dampens dimer formation (4). Although these interactions could facilitate the correct orientations for hetero-dimer assembly, such non-specific interactions could also potentially lead to inclusion bodies in red cells. The later is especially likely if the subunit concentrations are unequal and result in homo-oligomers (e.g. α_n_ or β_n_ occurring during thalassemia diseases) due to disrupted expression or genetic instability of either α or β subunits (20). *In vivo* experiments are needed to investigate how these non-specific interactions and molecular crowding influence the α_1_β_1_ dimer formation during Hb biosynthesis.

Finally, hemichromes are often observed within inclusion or Heinz bodies in red cells (12, 17). These particulates are found in red cells for hemoglobinopathies associated with globin instability or oxidative stress. The results in Figs. 6A and 6B provide an explanation for the presence of these species. At the high internal hemoglobin concentration in red cells, the unfolded holoprotein intermediates are dominant, leading to a much higher probability of misassembled Hb being trapped in the large hemichrome populations associated with the UH or IH_2_ states.

Our model was developed to examine unfolding as function of GdnHCl denaturant concentrations but could be modified to include other environmental or physiological Hb disassembly triggers such as low pH or high temperatures. Additionally, we can use the population fraction values extrapolated to 0 M GdnHCl to estimate the relative absolute free energies of different folding species *in vivo*. These relative free energies are defined as -RTlnY_i_ for species i, where a large positive value indicates a very unstable state (Fig. 8). This value estimates the amount of free energy that has to be added to disassemble or unfold the globin to a particular state. Fig. 8 shows that the free energy differences between apo- and holo intermediates are predicted to be higher in red cells compared to dilute plasma environments where hemin and subunit dissociation is promoted. However, in both environments the holo- and semi-holo disassembly intermediates are generally favored over the apo variants.

**Figure 8.**
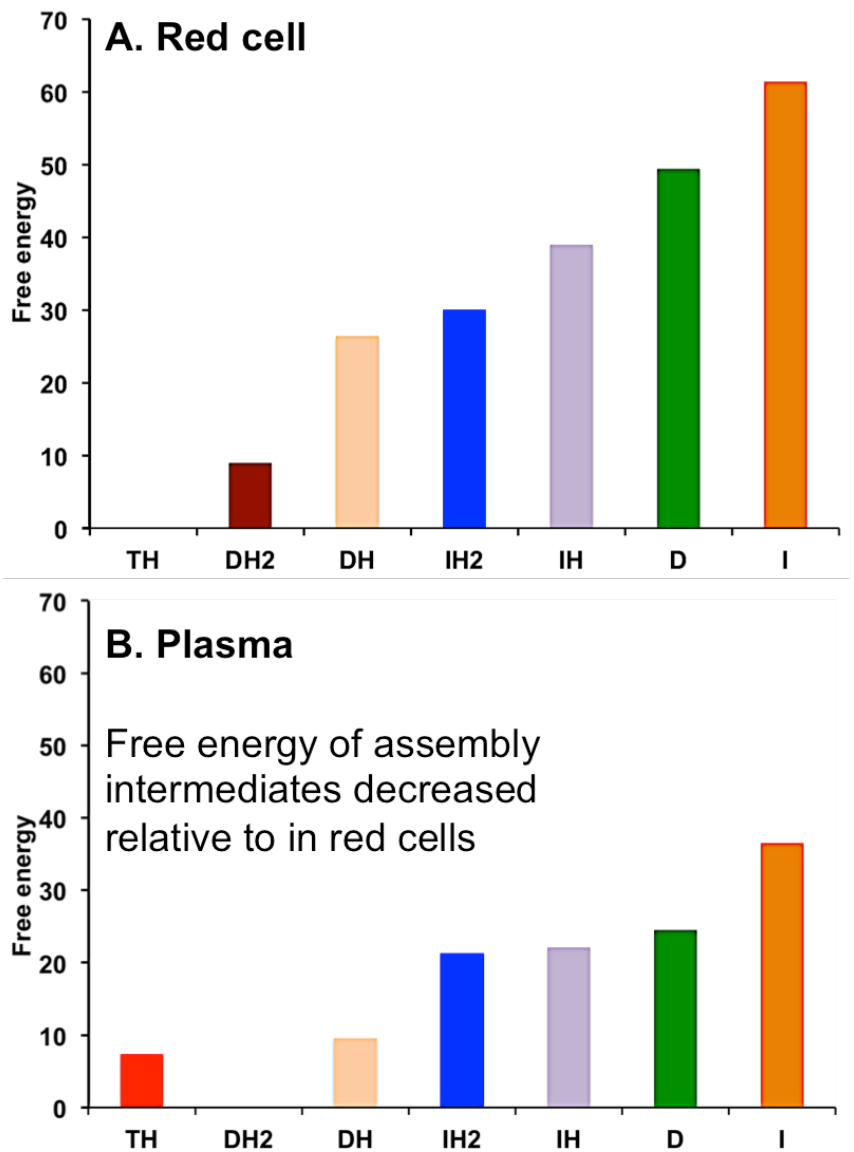
Predicted relative free energies of various HbA disassembly intermediates. In A. red cells and B. diluted into plasma.

When not contained within the reductive environment of red cells, acellular hemoglobin in plasma undergoes rapid autooxidation forming metHb, which in turn enables rapid heme disassociation. NO, which is generated in endothelial cells to facilitate vasodilation, can also react rapidly with acellular oxyHb to produce metHb and nitrate. A surge of acellular Hb in the bloodstream can lead to vasoconstriction due to NO scavenging by this dioxygenation reaction and subsequent release of hemin from acellular metHbA (82, 98–102).

The early onset of a significant population of disassociated free hemin in the low protein concentration pathway (Figs. 6C and 6D) provides an explanation for why chronic or severe hemolysis triggers free hemin mediated inflammatory diseases and endothelial dysfunctions that lead to serious cardiovascular complications and kidney problems (80, 103, 104). The disassociated free hemin intercalates into cell membranes and leads to the generation of reactive oxygen species (ROS) that can damage lipid, protein, and DNA (98, 105). Free hemin can also bind specifically to the toll-like receptor-4 (TLR4) on endothelial cells, triggering adverse pro-inflammatory cell cascades (106, 107). Hemin-mediated ROS generation and oxidative stress on the blood vessel walls also promotes adhesion molecule expression on the walls, leading to platelet aggregation and thrombosis (80, 98).

## Conclusion

We have been able to derive for the first time a quantitative Hb disassembly/assembly model, which can be used as a framework to identify the factors that resist Hb denaturation and those that enhance expression. The key feature of our model involves formation of molten globule intermediates with hemichrome spectral properties, and these intermediates are part of the pathway for the disassembly and unfolding of native hemoglobin. Complementary computational studies are also being done in our group to expand this framework to include interpretations at an atomic level. Hopefully, our efforts to obtain a comprehensive mechanism of Hb disassembly will lead to strategies for addressing erythropoiesis disorders and hemoglobinopathies in a clinical setting and for engineering more stable acellular hemoglobin-based oxygen carriers for transfusion therapy and other commercial purposes.

## Author contributions

PPS helped design the study, performed all the experiments, analyzed all of the data, and wrote all the drafts of the paper. The experiments were part of her Ph.D. thesis at Rice University under the guidance of JSO and GNP. She finalized the holoHb disassembly model and analyses at Rutgers University with the support of DAC. MAW performed and analyzed the SAXS experiments with samples provided by PPS. WCO expressed and purified the HbF samples and performed several HbF unfolding experiments as part of his undergraduate thesis at Rice University. GNP helped guide the work and edit various drafts of the paper. DAC also edited the paper and focused the modeling work on medical and physiological relevance. JSO helped to design the in vitro folding experiments, provided initial ideas about the model and analyses, and was the major editor of the manuscript.

## Acknowledgements

The research reported in this paper was funded by National Institutes of Health Grants HL110900 (JSO) and GM109456 (GNP), the BioXFEL Science and Technology Center Grant NSF 1231306 (GNP, DAC and PPS), Grant C-0612 from the Robert A. Welch Foundation (JSO and PPS), and the Sealy Center for Structural Biology and Molecular Biophysics at UTMB. We would like to thank Thomas Grant for reviewing our SAXS data and analyses and Jayashree Soman and Eileen Singleton for help in expressing, purifying, and running quality control analyses of many of the recombinant hemoglobins.

## Supplemental Information

Additional supplementary figures and a table are provided.

