## Supplemental figures and table for "The interplay between molten globules and heme disassociation defines human hemoglobin disassembly"

### Supplementary Figures and Table

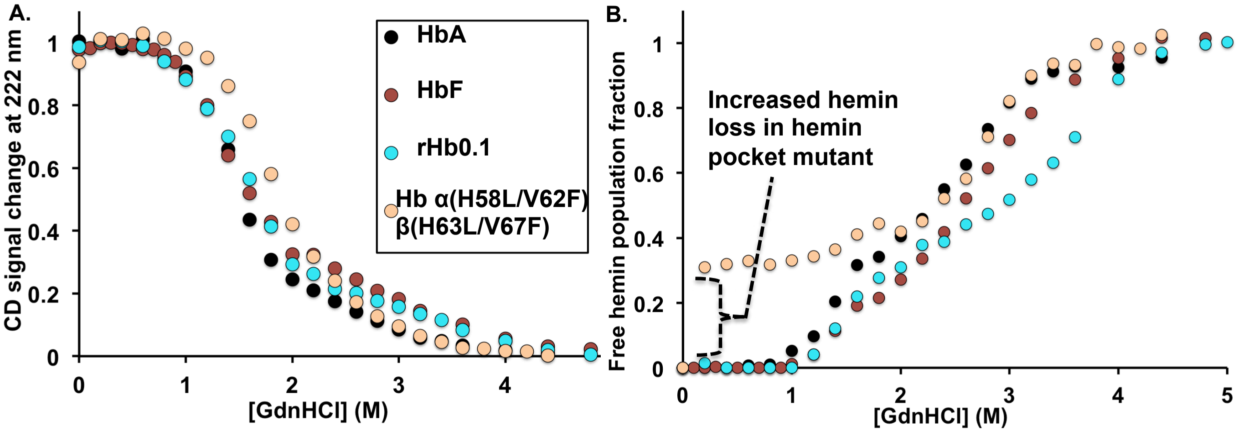


Figure supplement 1. **Comparison of GdnHCl-induced equilibrium unfolding measurements of holo- HbA HbF, rHb0.1, and Hb α(H58L/V62F)β(H63L/V67F).** The solid circles are the measured data at 12 μM per subunit total protein concentration. ***A,*** Fractional CD change at 222 nm, a measure in the change of alpha-helical structure content relative to folded holoHb. ***B,*** Population fractions of free hemin. The experimental measurements in panel B were obtained from deconvolution of visible absorbance spectra recorded between 350 nm and 660 nm. All holoHb unfolding measurements were done in 200 mM potassium phosphate, pH 7 at 10 °C. Protein samples were initially prepared in met oxidation states. The 12 μM HbF CD unfolding measurement was obtained from previously published data [4].

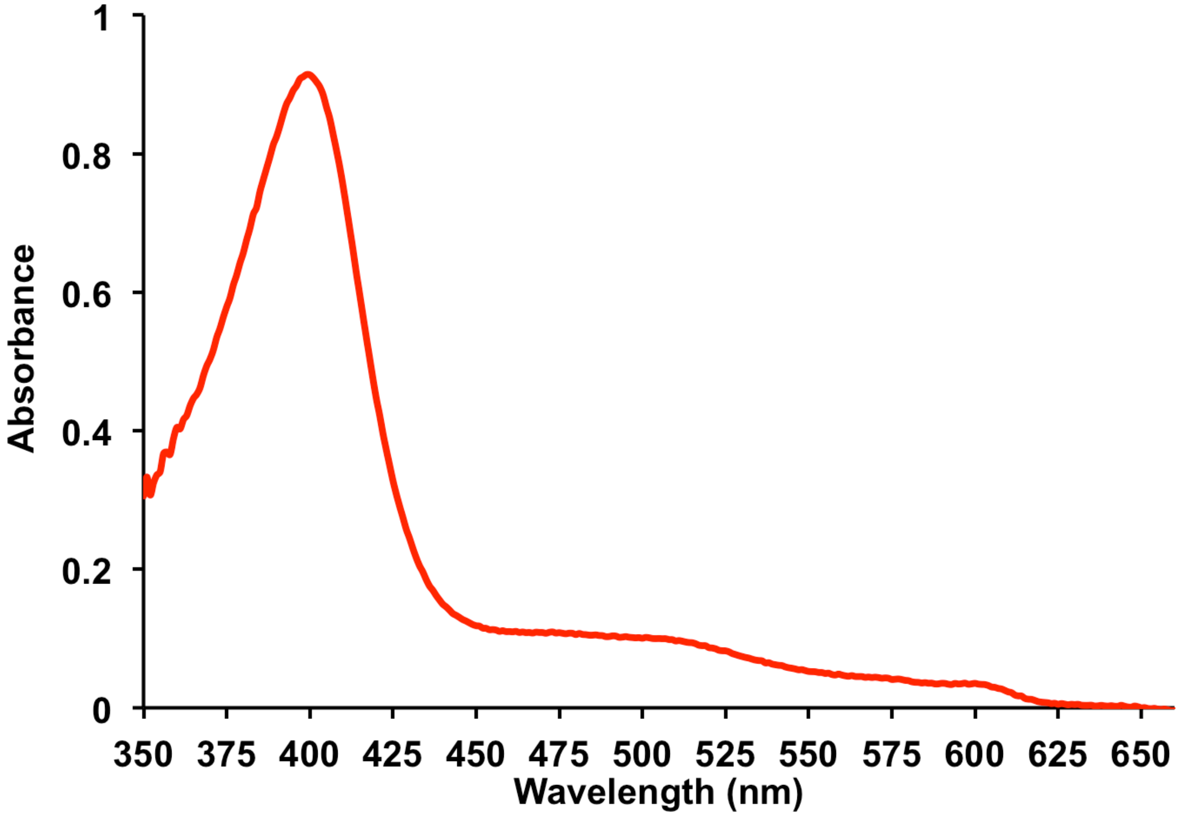


Figure supplement 2. **Visible absorbance spectra for Hb α(H58L/V62F)β(H63L/V67F)** at 12 μM per subunit protein concentration. Measurements were done in 200 mM potassium phosphate at pH 7.

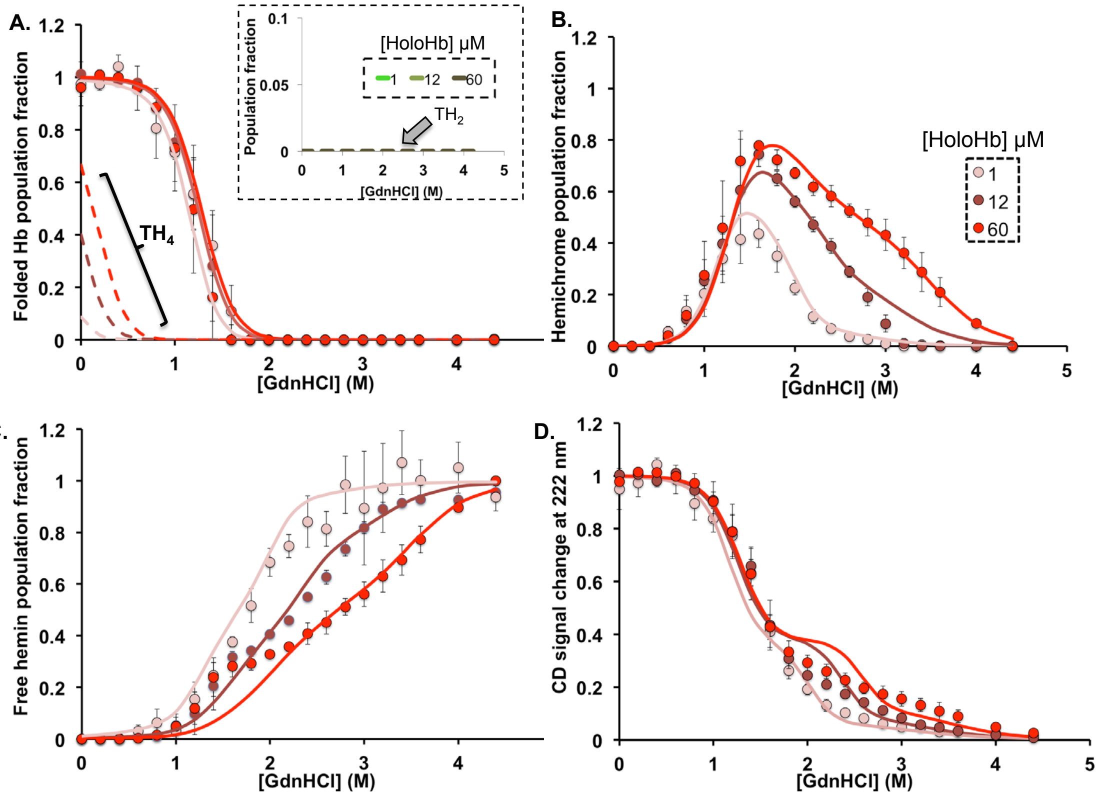


Figure supplement 3. **Fitting of GdnHCl-induced disassembly measurements for HbA to equilibrium disassembly model 2**. The solid circles are the measured data at different per subunit total protein concentration and the corresponding solid lines are the fits. ***A,*** Population fractions of aquamet- holoHb dimers and tetramers. The dashed lines outset and inset are the theoretical population fractions of folded holoHb tetramers ($TH_4$) and semi- holoHb tetramers ($TH_2$), respectively, derived from the fits. ***B,*** Population fractions of hemichromes. ***C,*** Population fractions of free hemin. ***D,*** Fractional CD change at 222 nm, a measure in the change of $\alpha$-helical content relative to folded holoHbA. The experimental measurements in panels A-C were obtained from deconvolution of visible absorbance spectra recorded between 350 nm and 660 nm. All holoHb unfolding measurements were done in 200 mM potassium phosphate, pH 7 at 10 °C. Protein samples were initially prepared in aqua-met oxidation states.

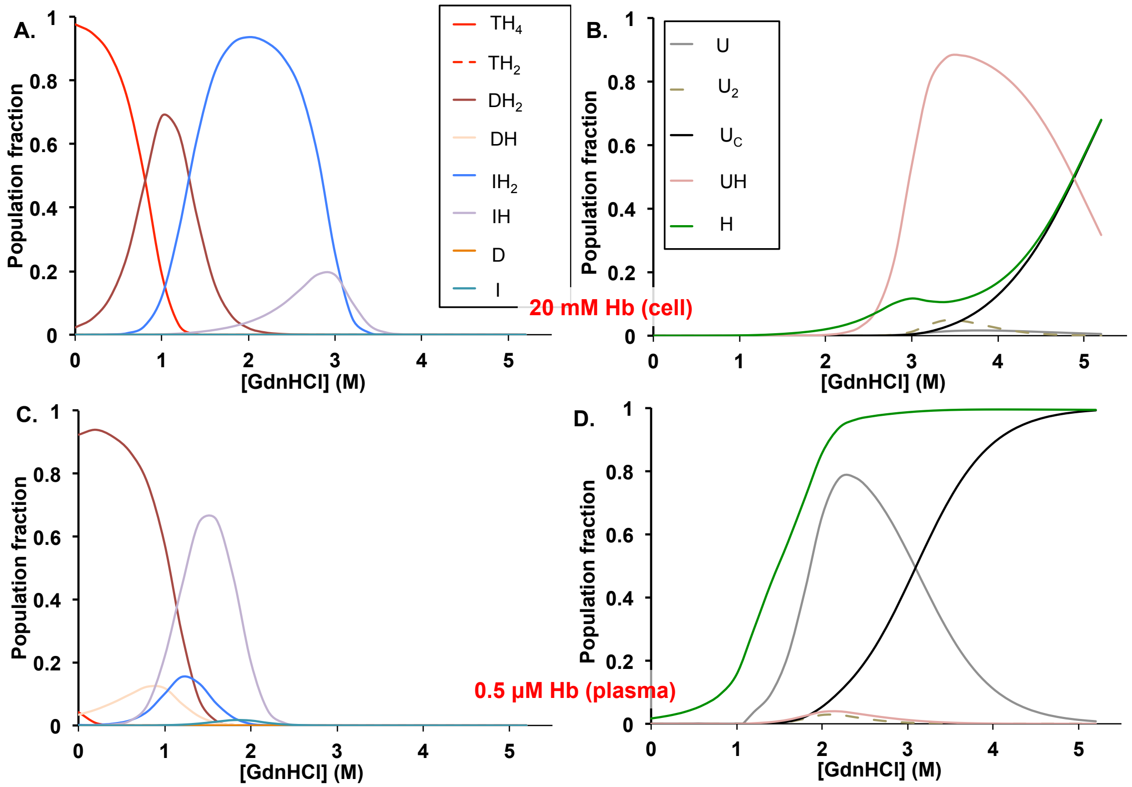


Figure supplement 4. **Simulated HbA disassembly population fractions** in ***A*****, Hb packed in red blood cells** and ***B*****, Hb diluted in plasma** using Model 2 (blue and black arrows in Figure 1). Simulations were done with parameters from Table S1.

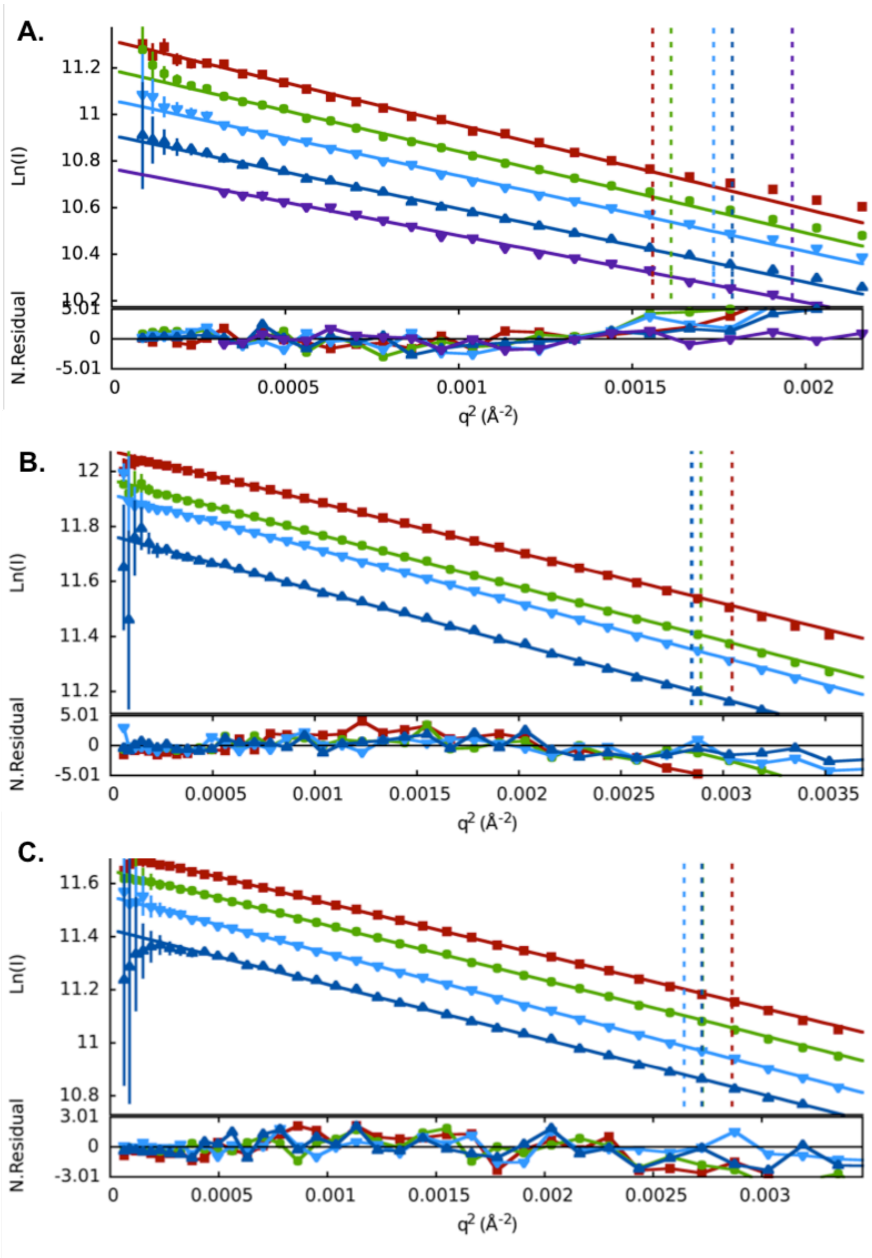


Figure supplement 5. **The Guinier Plots of each Hb dilution series.** ***A,*** The aporHb0.1 dilution series (0.0 (extrapolated),1.5, 2.0, 2.5, and 4.0 mg/ml). ***B,*** The holorHb0.0 dilution series (1.0, 2.0, 3.0, and 6.0 mg/ml). ***C***, The holorHb0.1 dilution series (1.0, 2.0, 3.0, and 6.0 mg/ml). Curves are heatmap colored by concentration and offset for clarity. The dashed vertical lines mark the upper q-limits of the Guinier fits ($1.3/R_g$). The aporHb0.1 series shows a positive concentration dependence indicative of molecular crowding at protein concentrations >1.5mg/ml. The two holo-Hb series display a negative concentration dependence indicative of electrostatic repulsion in the 6 mg/ml samples.

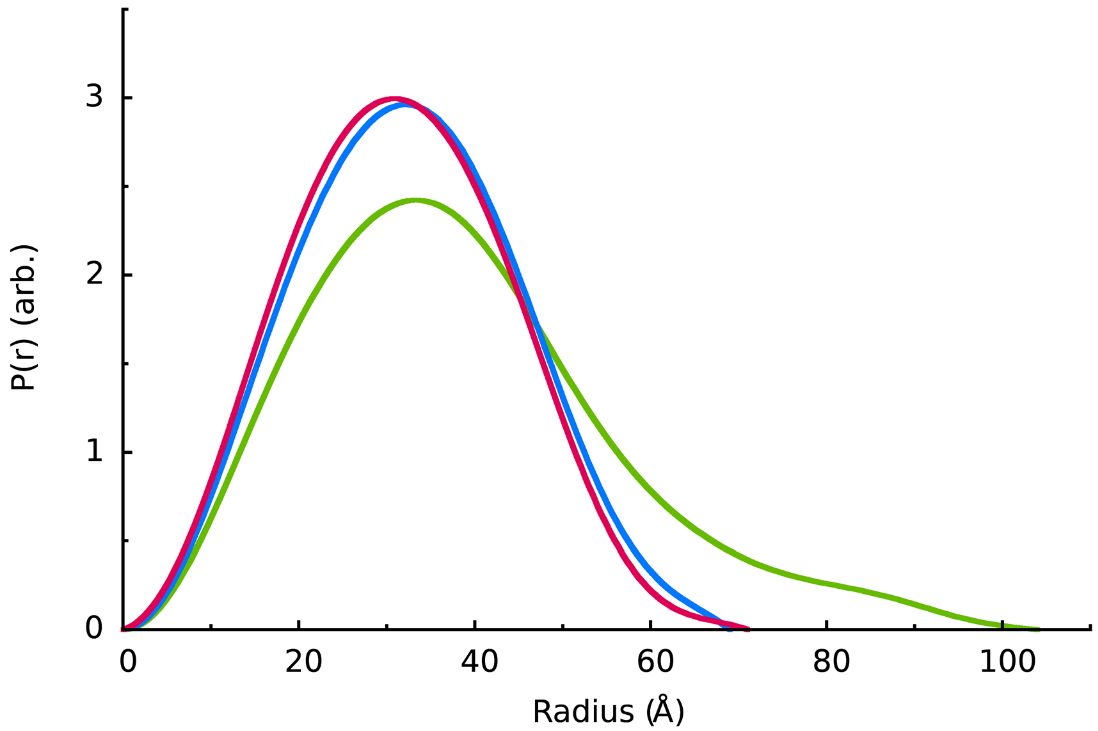


Figure supplement 6. **The GNOM P(r) plots** for the zero-concentration extrapolated **aporHb0.1** (green), 2.0 mg/ml **holorHb0.1** (blue), and 2.0 mg/ml **holorHb0.0** (red). From the curves, Dmax values for aporHb0.1: 104 Å, holorHb0.1: 69 Å, holorHb0.0: 71 Å. The area under each curve was normalized to the same arbitrary value, the area for holorHb0.0. The broadening of the distribution curve for aporHb0.1 suggests that it has become expanded relative to more compact holoHb forms.

Table supplement 1. **Fitted equilibrium disassembly parameters to model 2 for HbA**

| | HbA |
|---|---|
| $K_{2,4}$ ($M^{-1}$) | $1.0 \times 10^{5}$ |
| $K_{DH,DH2}$ ($M^{-1}$) | $3.3 \times 10^{9}$ |
| $K_{D,DH}$ ($M^{-1}$) | $5.2 \times 10^{11}$ |
| $K_{IH,IH2}$ ($M^{-1}$) | $6.0 \times 10^{8}$ |
| $K_{I,IH}$ ($M^{-1}$) | $3.1 \times 10^{11}$ |
| $K_{U,UH}$ ($M^{-1}$) | $1.2 \times 10^{6}$ |
| $K_{TH4,TH2}$ ($M^{2}$) (per 2 subunits) | $1.7 \times 10^{-25}$ |
| $m_{2,4}$ (kJ $mol^{-1}$ $M^{-1}$) | 20.0 |
| $m_{DH,DH2}$ (kJ $mol^{-1}$ $M^{-1}$) | 9.3 |
| $m_{D,DH}$ (kJ $mol^{-1}$ $M^{-1}$) | 9.3 |
| $m_{IH,IH2}$ (kJ $mol^{-1}$ $M^{-1}$) | 10.9 |
| $m_{I,IH}$ (kJ $mol^{-1}$ $M^{-1}$) | 10.9 |
| $m_{U,UH}$ (kJ $mol^{-1}$ $M^{-1}$) | 2.5 |
| $m_{TH4,TH2}$ (kJ $mol^{-1}$ $M^{-1}$) | 9.3 |
| $S_{TH4}$, $S_{TH2}$ | 1.00, 0.87 |
| $S_{DH2}$ | 1.00 |
| $S_{DH}$ | 0.87 |
| $S_{IH2}$ | 0.35 |
| $S_{IH}$ | 0.42 |
| $S_{UH}$ | 0.02 |
